# High-throughput, fluorescent-aptamer-based measurements of steady-state transcription rates for *Mycobacterium tuberculosis* RNA polymerase

**DOI:** 10.1101/2023.03.13.532464

**Authors:** Drake Jensen, Ana Ruiz Manzano, Maxwell Rector, Eric J. Tomko, M. Thomas Record, Eric A. Galburt

## Abstract

The first step in gene expression is the transcription of DNA sequences into RNA. Regulation at the level of transcription leads to changes in steady-state concentrations of RNA transcripts, affecting the flux of downstream functions and ultimately cellular phenotypes. Changes in transcript levels are routinely followed in cellular contexts via genome-wide sequencing techniques. However, *in vitro* mechanistic studies of transcription have lagged with respect to throughput. Here, we describe the use of a real-time, fluorescent-aptamer-based method to quantitate steady-state transcription rates of the *Mycobacterium tuberculosis* RNA polymerase. We present clear controls to show that the assay specifically reports on promoter-dependent, full-length RNA transcription rates that are in good agreement with the kinetics determined by gel-resolved, α-^32^P NTP incorporation experiments. We illustrate how the time-dependent changes in fluorescence can be used to measure regulatory effects of nucleotide concentrations and identity, RNAP and DNA concentrations, transcription factors, and antibiotics. Our data showcase the ability to easily perform hundreds of parallel steady-state measurements across varying conditions with high precision and reproducibility to facilitate the study of the molecular mechanisms of bacterial transcription.

**Significance Statement:** RNA polymerase transcription mechanisms have largely been determined from *in vitro* kinetic and structural biology methods. In contrast to the limited throughput of these approaches, *in vivo* RNA sequencing provides genome-wide measurements but lacks the ability to dissect direct biochemical from indirect genetic mechanisms. Here, we present a method that bridges this gap, permitting high-throughput fluorescence-based measurements of *in vitro* steady-state transcription kinetics. We illustrate how an RNA-aptamer-based detection system can be used to generate quantitative information on direct mechanisms of transcriptional regulation and discuss the far-reaching implications for future applications.

## Introduction

Cellular RNA abundance is dictated by the relative steady-state rates of RNA production and degradation. In particular, the rate of RNA production is ubiquitously the target of gene regulatory mechanisms and often represents a good proxy for protein synthesis flux (1). More specifically, the rate at which full-length RNA transcripts are generated is typically controlled by the rate of transcription initiation. This is because the overall initiation rate is often slower than subsequent elongation and termination steps, and because multiple RNA polymerases (RNAPs) may be elongating the same template at the same time. Bacterial transcription initiation progresses through several intermediates, where the rates and equilibrium constants that describe the initial binding of the RNAP to the promoter and the subsequent isomerization steps that culminate in opening of promoter DNA can vary greatly depending on the sequence of the promoter (2, 3). For example, *E. coli* promoters can differ in rates of promoter opening by factors of 10^3^ – 10^4^, resulting in initiation events ranging from once per second to once per generation (4, 5). Following promoter opening, binding of the first two initiating nucleoside triphosphates (NTP) forms an initially transcribing complex that begins producing a nascent RNA transcript. Translocation events accompanying early synthesis of short RNA transcripts are accompanied by disruption of RNAP-promoter contacts, slowing these steps. Once all contacts with the promoter are broken, nucleotide incorporation kinetics become faster and the RNAP enters the processive elongation phase of transcription, escaping the promoter. After decades of careful biophysical dissection, driven mainly by pre-steady-state kinetic and structural biology approaches, bacterial initiation pathways are well-characterized on a handful of different promoters for model bacteria (reviewed in (2, 3, 6–8)). Many kinetic/structural intermediates have been identified, including off-pathway states that, in some cases, do not lead to full-length RNA production (9–14). The resulting models from all these studies are complex and can vary depending on the system studied (15–17), bringing into question model generalizability across bacteria and different promoter sequences.

Ideally, rate constants obtained from pre-steady-state kinetic measurements could be used to calculate an overall initiation rate or the average time between initiation events, i.e., the time required for RNAP binding, DNA opening, initial transcription, and promoter escape. However, while simplified models may address theoretical links between initiation kinetics and steady-state rates of RNA production (3, 18), they cannot quantitatively predict how changes in promoter sequence and regulatory conditions affect steady-state flux. This overall initiation rate is most simply described using Michaelis-Menten enzyme kinetics, where the RNA polymerase is the enzyme, promoter DNA is the substrate, and the full-length RNA transcript is the product (19). This model assumes a constant concentration of RNAP–promoter (enzyme–substrate) complexes generated by a balance between promoter binding and escape rates and predicts a constant reaction velocity or rate of transcript production. Within this formalism, regulated promoters are activated or repressed via changes in the Michaelis-Menten parameters *K*_*m*_ (the concentration at which the half maximal rate is observed) and/or *V*_*max*_ (the maximal rate observed under saturating conditions) without changes in the free RNAP concentration. Alternatively, constitutively active promoters are affected by cell growth-rate-dependent changes in the free RNAP concentration, independent of changes in *K*_*m*_ and/or *V*_*max*_ (20–22). Classic examples of environmental adaptation in bacteria affecting transcription initiation processes include the up-regulation of beta-galactosidase in response to the presence of lactose and the absence of glucose (23), and the genome-wide response to starvation known as the stringent response (24, 25).

*In vitro* measurements of steady-state transcription rates can be used to test and develop more complex models but have been limited by current methodologies. Historically, measurements of *in vitro* basal and regulated transcription initiation kinetics have been made possible by monitoring RNA production via the incorporation of radio-labeled NTPs (26). This assay affords high sensitivity and single-nucleotide resolution even with small quantities of RNA when resolved by gel electrophoresis. This experimental approach was used in the initial discovery of RNAP holoenzyme activity more than fifty years ago (27) and is still the most common assay for probing transcriptional mechanisms *in vitro*. However, steady-state ^32^P-detected transcription assays have several drawbacks. A practical limitation is that the use of radioactivity is expensive, both for the purchase of the reagent itself and for the requirement of specialized protocols for safe shipping and use in the lab. The reaction must also be taken through several steps that may include the incorporation of proteases or chelators to quench the reaction at set time points, phenol/chloroform extractions, and/or denaturation and loading the sample on an electrophoretic polyacrylamide gel. All these steps, as well as the quantification of resolved product bands via image analysis, can introduce non-biochemical variability in the measured RNA amounts. In addition, these assays require significant amounts of time and training, where single experiments often take multiple days before complete quantification and where technical practice is needed for generating reproducible data. All these factors combined cause ^32^P detection assays to have relatively limited throughput. As a result, the time-dependent measurements needed to quantitate steady-state initiation rates are infrequently performed and single time-point measurements are often used to infer mechanisms of gene regulation.

In contrast to the *in vitro* approaches described above, *in vivo* transcription studies have undergone a remarkable transformation where genome-wide transcript levels in cells under varying environmental and genetic backgrounds are routinely queried (28). We were motivated by the desire to increase the throughput of *in vitro* transcriptional studies to facilitate the efficient testing of mechanistic hypotheses, the development of predictive transcription initiation models, and to be able to compare transcript flux from reconstituted systems to the genome-wide information accessible via RNAseq. To this end, we explored the use of a fluorescent-aptamer-based detection system to report on *in vitro* steady-state transcription rates.

Fluorescent light-up aptamers are RNA sequences that bind small molecule fluorophore dyes and generate a large fluorescence enhancement. They have been used in numerous applications (reviewed in (29)) including the detection of nascent RNA transcripts in cells (30–35), in synthetic biology transcription-translation coupled *in vitro* systems for T7 RNAP (36–38) and with cell lysates from diverse bacterial species (39). These approaches rely on the use of a DNA template that encodes for an aptamer sequence so that each transcription event results in the generation of an RNA aptamer which folds and binds the dye. For each transcript produced, there is an accompanying increase in fluorescence (**Figure 1**). Importantly, as the change in dye fluorescence requires the synthesis of a full-length RNA transcript containing the aptamer, the fluorescent readout is not complicated from short abortive products that may be generated during promoter escape. Using this approach, recent work has illustrated how the rates of *in vitro* reactions can be quantified in a fluorimeter cuvette with *E. coli* RNAP (40) and in a plate-reader format with T7 RNAP (41). Here, we follow up on these studies and provide a description of essential control experiments needed to clearly link the fluorescent signal with the transcription of a promoter-derived product. Additionally, using data acquired with RNAP from *Mycobacterium tuberculosis* (*Mtb*), we describe a workflow for the calibration and quantification of RNA concentration in multi-round initiation kinetics analyzed with a Michaelis-Menten approach. Most significantly, we illustrate the utility of this approach for the investigation of mechanisms of transcriptional regulation by NTP concentration, transcription factors, and antibiotic inhibitors, all derived from high-throughput measurements under steady-state conditions.

**Figure 1.**
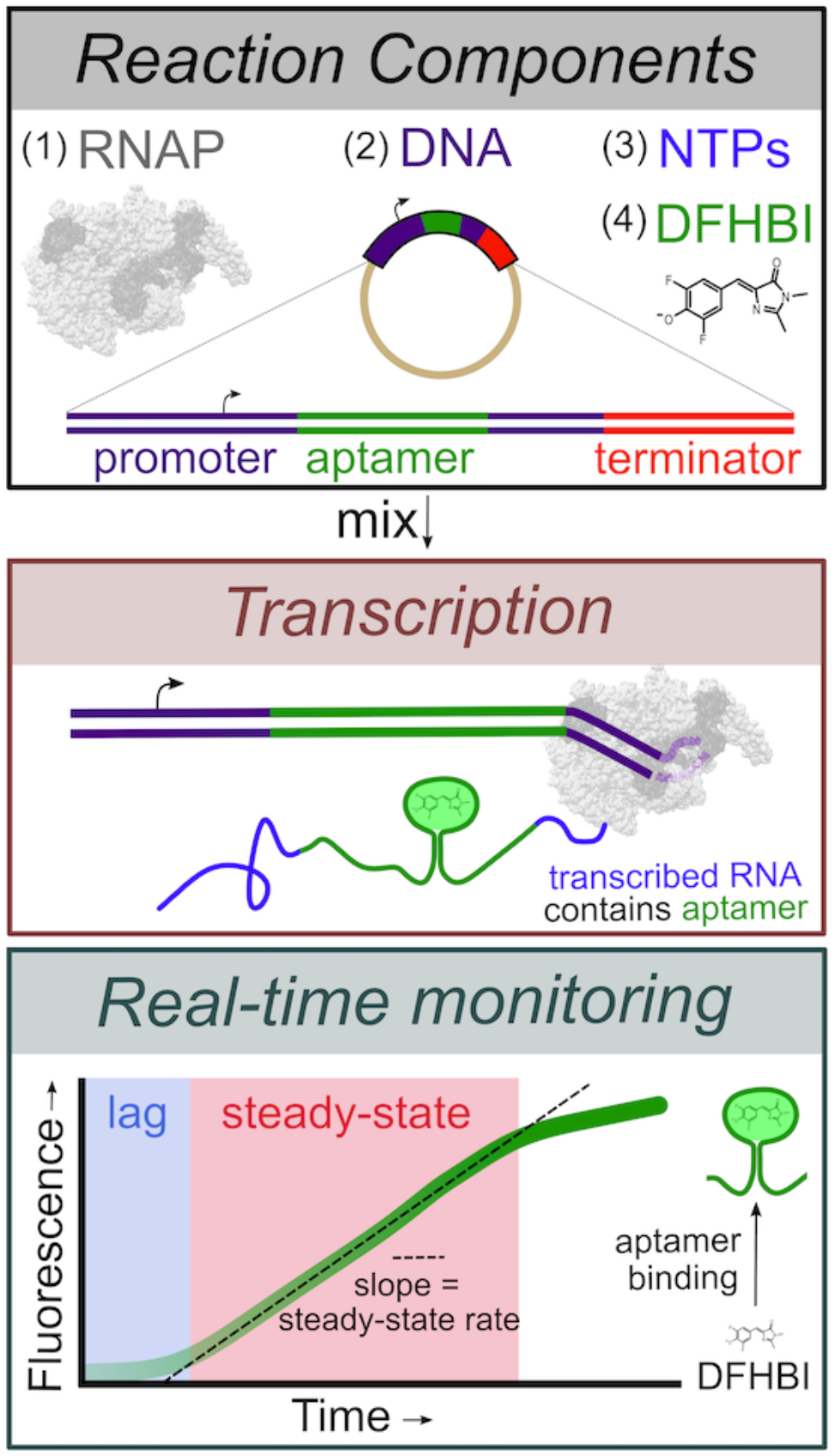
Overview of the fluorescent-aptamer-based assay. This assay requires RNAP, a DNA template containing the sequence for the Spinach-mini RNA aptamer (30), NTPs and the fluorescent dye DFHBI. Upon initiating the reaction, full-length RNAs containing the Spinach-mini aptamer capable of binding DFHBI are synthesized. The fluorescent signal change, monitored in real-time, results from the unbound to aptamer-bound dye transition and is used as a reporter of full-length transcription rates. An initial lag time is observed, followed by a steady-state regime where the slope of the linear phase represents the steady-state rate of full-length RNA synthesis.

## Results

### Promoter-dependent, aptamer-based measurements of full-length transcription

As similar assays have been described elsewhere, we briefly describe the specifics of the version we utilized for these studies. DNA templates were constructed such that an 80 nt Spinach-mini aptamer sequence (30) was inserted 30 bp downstream of the +1 transcription start site corresponding to the genomic *Mtb* ribosomal RNA promoter sequence (*rrnA*P3). Specifically, –60 to +31 of *rrnA*P3 was included to account for any upstream-promoter interactions as well as the initially transcribed sequences, both of which can regulate initiation kinetics. Inclusion of the genomic initially transcribed sequence, rather than just positioning the aptamer sequence to start at the +1 position, provides for unaltered promoter escape, which is known to become rate limiting under certain conditions (see (3) for examples of sequence effects on initiation kinetics). The *E. coli* intrinsic *rrnB*P1 *T*_1_ terminator sequence was included in DNA plasmid templates to dissociate the ternary polymerase, template, and transcript complex. Sequences for both plasmid and linear templates are given in **Supplementary Tables 1 and 2**.

When transcribed downstream of the *Mtb rrnA*P3 promoter, the Spinach-mini aptamer sequence folds and binds the small molecule fluorophore DFHBI, resulting in a fluorescence enhancement. By mixing RNAP, DNA, all NTPs, and DFHBI, the reaction is monitored in real-time where the increase in fluorescence reports on transcript production. All experiments were performed in a 384 well plate-reader format using 10 µL reaction volumes facilitating high-throughput measurements of steady-state initiation kinetics with minimal sample volume requirements. The slope of the fluorescent signal at early times reports on the steady-state rate of transcription initiation (**Figure 1**). In general, one expects initiation kinetics (on the order of minutes to hours) and not co-transcriptional aptamer folding (estimated to be on the order of seconds to minutes (36, 42, 43)) or dye binding (44) to be rate-limiting for functional aptamer production. However, even when the initiation rate is not limiting relative to the timescales of elongation or aptamer folding, the steady-state rate will still specifically report on the initiation rate (**Supplementary Figure 1**). This is because once one RNAP leaves the promoter, another RNAP may bind and begin the process of initiation irrespective of these downstream processes. This condition may be broken by pathological cases where downstream pausing creates a traffic jam that backs up onto the promoter, influencing the time needed for the next polymerase to bind the promoter.

Transcription reactions were initiated by adding DNA templates to *Mtb* RNAP preincubated with NTPs, DFHBI dye, and RNase inhibitors. Fluorescence was monitored in real-time, where we observed three distinct kinetic phases to the trace using the plasmid template containing the *rrnA*P3 sequence: 1) a lag time, 2) a linear increase, and 3) roll-off from the linear regime that begins to plateau over time (**Figure 2A**, black). We note that when starting from a well-controlled time zero, the lag time theoretically reports on the time required for pioneering RNAPs to complete the initiation process, transcribe the aptamer, and for dye to bind the folded RNA transcript (40) (**Supplementary Figure 1**). In our assay setup, since initiation of the reaction was done by hand, we do not attempt to quantitate the lag time, but focus on the linear regime. In the absence of either DNA or NTPs, only a slow decay in DFHBI fluorescence was observed as a function of time (**Supplementary Figure 2**), indicating that the increase in fluorescence when all reaction components are present is due to transcription-derived aptamer formation. For all plots shown, the signal in the absence of transcription is subtracted from the experimental traces so that the y-axis reports specifically on the fluorescence generated by transcription of the aptamer (Methods). Correcting for this time-dependent fluorescent change of the background signal is especially important when evaluating conditions of minimal transcript generation (**Supplementary Figure 2**).

**Figure 2.**
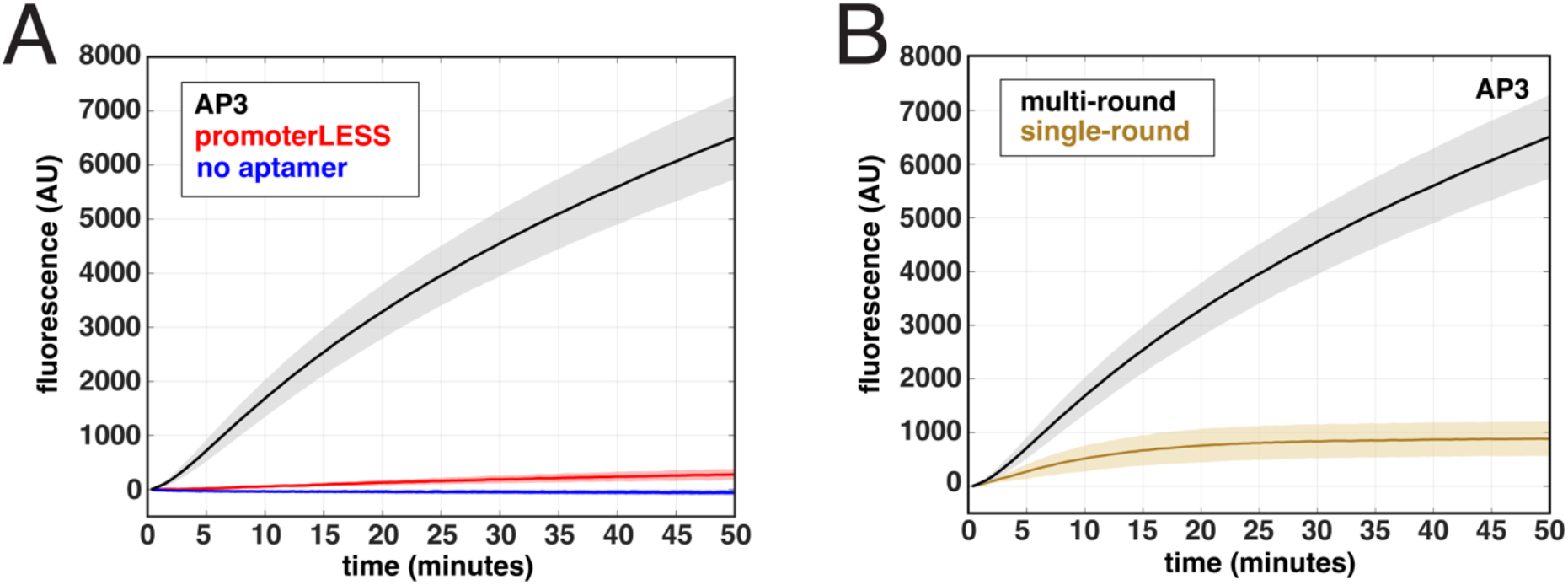
Real-time fluorescent signal is promoter dependent and due to multiple rounds of initiation. All experiments were performed using 100 nM *Mtb* RNAP, 5 nM DNA plasmid templates, and 500 μM NTPs. **(A)** Comparison of time courses of RNA synthesis from an *Mtb* plasmid template containing either both the *rrnA*P3 and aptamer sequences (black), the *rrnA*P3 sequence with no aptamer sequence (blue), or the “promoterLESS” template, containing the aptamer, but lacking the *rrnA*P3 sequence (red). **(B)** Real-time traces comparing multi-round (black) and single-round conditions (gold) on the *rrnA*P3 template. Error shading indicates the standard deviation of three biological replicates.

To confirm that transcription is driven by the *rrnA*P3 promoter sequence, we performed the assay using a plasmid template lacking the promoter (i.e., “promoterLESS”; for sequence, see **Supplementary Table 1**) as a negative control. The signal generated in the presence of *rrnA*P3 is more than 20-fold higher than that obtained with the promoterLESS template (**Figure 2A**, red), confirming that the signal arises from promoter-driven transcription. However, the amount of signal generated by the promoterLESS template is not zero, as can be seen by comparing to a trace using a template that contains the promoter but lacks the aptamer sequence (**Figure 2A**, blue). This comparison suggests that even in the absence of a promoter sequence, some basal level of aptamer is transcribed (see below for further discussion).

### Fluorescence can be monitored under both multi- and single-round conditions

Our specific goal was to obtain measurements of steady-state rates of transcription. We therefore needed to confirm that the observed signal is due to multiple rounds of transcription. We expect significantly higher signal amplitude under multi-round conditions compared to single-round conditions, where DNA traps are used to prevent RNAP rebinding. Use of salmon-sperm DNA, which does not actively dissociate RNAPs from the DNA like other competitors such as heparin (17), established a baseline amplitude (Methods) (**Supplementary Figure 3A**), serving as an effective trap for the *Mtb* RNAP. Upon initiating the transcription of pre-bound RNAP-DNA complexes with 500 μM NTPs and salmon-sperm DNA competitor, we observed no lag time, and the fluorescence rapidly reached saturation as RNAP re-binding was prevented (**Figure 2B**). These results are consistent with single-round conditions of full-length transcript production measured under our detection times (10, 13, 45). In the absence of competitor under otherwise identical conditions, we observed a large increase in the overall fluorescent signal (**Figure 2B**), confirming that in the absence of DNA trap, the assay is multi-round.

In theory, use of this assay under single-round conditions with pre-formed RNAP-DNA complexes should permit kinetic analyses of process that are difficult to determine in multi-round conditions, such as promoter escape. Titrating all four NTPs in the presence of the DNA trap resulted in an NTP-concentration-dependent increase in signal amplitude, where the traces could be well-fit by a single-exponential function (**Supplementary Figure 3B**). This NTP-concentration-dependent change in amplitude suggests that the rate of RNAP dissociation from the promoter is on the same order as the rate of promoter escape under these conditions. This observation is consistent with our previous transient-state kinetic measurements of *Mtb* promoter escape kinetics, where we observed that increasing NTP concentration and incorporation of the initially transcribed sequence stabilizes the RNAP-DNA complexes, slowing the rate of dissociation and facilitating escape (17). These results illustrate that this fluorescent-aptamer-based assay can be used under single-round conditions to examine the kinetics of sub-steps in initiation (i.e., dissociation and promoter escape).

### Extracting steady-state rates from real-time, multi-round fluorescent traces

Under multi-round conditions to examine the steady-state rates of transcription, we observed that the fluorescent time traces exhibited an apparent lag, followed by a linear increase in fluorescence over time. This rate of increase slows over longer timescales and eventually begins to plateau (**Figure 3A**, solid lines). We hypothesize that saturation of the fluorescence signal can be explained by the presence of paused and/or backtracked polymerases trapped on the template (reviewed in (46)), leading to a reduction in the RNAP molecules that can re-initiate at the promoter over time. Consistent with this hypothesis, traces collected with *E. coli* RNAP in the presence and absence of the RNA cleavage factor GreB overlap at early time points and then diverged, where conditions containing GreB resulted in a higher fluorescent signal (**Supplementary Figure S4**). Based on thse results, we suggest that GreB acts to increase the active RNAP concentration at longer times by facilitating recycling of long paused/backtracked elongation complexes (47, 48). As GreB had no effect in the initial linear region, we suggest that this region reports on the true steady-state rate of initiation, without interference from inactive elongation states occurring downstream of the promoter. In addition, these results suggest that the fluorescent-aptamer-based assay can also be used to examine the regulation of rate-limiting elongation processes at long timescales.

**Figure 3:**
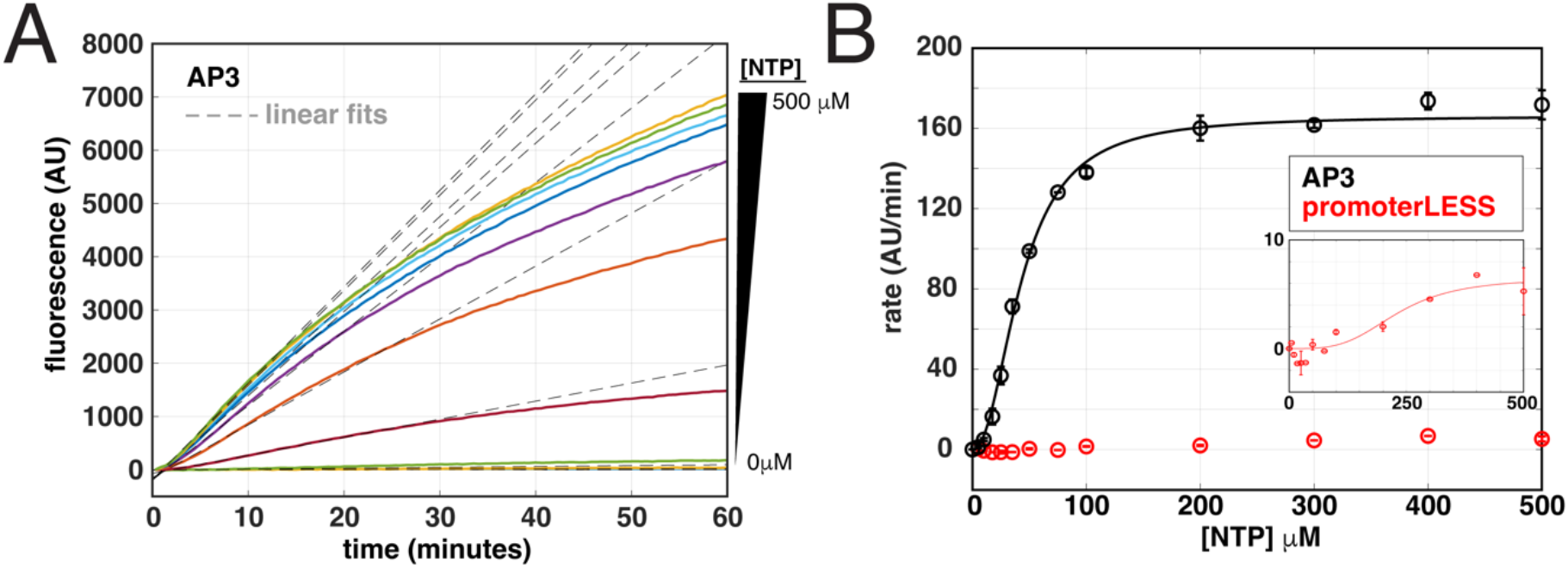
Quantification of the NTP-dependence of steady-state rates for full-length transcription. **(A)** Titration of the concentration of all NTPs with 100 nM *Mtb* RNAP and 5 nM *rrnA*P3 plasmid DNA results in an increase in the rate and amplitude of the fluorescent traces (solid, colored lines). The unbiased linear fits of the early times are shown in gray dotted lines for each trace. **(B)** Quantification of steady-state rates on *rrnA*P3 (black) and promoterLESS (red) DNA plasmid templates plotted as a function of NTP concentration and fit to a modified Michaelis-Menten model (**Eq. 2**) to account for the apparent sigmoidal behavior.

Based on these results, we analyzed the initial fluorescence increase to quantitate the steady-state rate. Previous work manually determined a time regime of the trace to fit with either a line or an exponential. The slope was then determined directly or through differentiation, incorporating an initial time offset to account for the lag-phase (40). Here, to define the steady-state (linear) regime, we designed an automated fitting protocol to reproducibility fit large amounts of data with minimum user input (Methods). Briefly, for each biological replicate, an average of two – three technical replicates were recursively fit with a line, using a statistical weight determined by the variation between technical replicates. The length of the time frame in each round of fitting is reduced in each cycle and the R^2^ value (goodness of fit) is compared to a threshold set by the user to evaluate whether the fit was adequately linear or whether a further reduction in the time regime should be attempted (**Supplementary Figure S5**). For each trace, the code then plots the raw data and the fit for visual inspection by the user. Finally, the estimated rates are outputted for subsequent analyses. We have made this code publically available on Github (https://github.com/egalburt/aptamer-flux-fitting).

### Steady-state kinetic measurements are dependent on NTP concentration

We examined the NTP concentration dependence of reaction rates to ensure that the time-dependent signal change in fluorescence follows our expectations of steady-state behavior. In addition to promoter DNA, NTPs can also be considered as a substrate in the Michaelis-Menten analysis of the transcription reaction. In this case, concentrations below those required to reach *V*_max_ can lead to a decrease in the rate of transcription, generally attributed to slower nucleotide incorporation rates and/or increases in the abortive fraction (10, 17). Titrating all four NTPs together up to 500 µM on the *rrnA*P3 plasmid template, we observed a clear concentration dependence in the real-time traces (**Figure 3A**, solid lines). We fit the entire data set with our variable-time fitting algorithm described above, permitting us to extract the steady-state rates at each NTP concentration (**Figure 3A**, dotted lines). We plotted these rates as a function of NTP concentration and fit the data to a modified Michaelis-Menten equation, weighted by the exponential parameter *n* (**Eq. 2**, Methods) (**Figure 3B**). The data did not fit the simpler form of the equation where *n* = 1 and an unconstrained fit results in an *n* = 2.1 ± 0.4. When *n* > 1, it signifies a steeper concentration dependence than expected from Michaelis-Menten equation for a single substrate binding site. Possible interpretations for this non-hyperbolic behavior can be found in the Discussion.

We performed analogous NTP titrations on the promoterLESS plasmid template. Compared to the kinetic parameters obtained with the *rrnA*P3 plasmid, we observed an ∼25-fold decrease in *V*_*max*_ (6.5 ± 3.8 AU/min compared to 165 ± 7 AU/min) and an ∼5-fold increase in the apparent *K*_*m*_ (*K*_*m,app*_, defined in Methods, 230 ± 120 μM compared to 44 ± 5 μM) (**Figure 3B**, inset). As mentioned above, given that the signal from the promoterLESS plasmid is above that of the no-aptamer control obtained at saturating NTPs (**Figure 2A**), we interpret the NTP-dependent signal using the promoterLESS template as a measure of the amount of aptamer transcription derived from non-specific RNAP-DNA promoter interactions. Being able to measure these NTP-concentration-dependent trends in signal demonstrates that the assay possesses the detection sensitivity to needed measure low concentrations of full-length transcripts, including those produced from sequences nominally devoid of promoters. As the promoterLESS control sets a lower bound for detecting promoter-dependent transcription, it should always be included and, if need be, used as a correction. In this specific case, subtraction of the promoterLESS signal from that obtained with *rrnA*P3 resulted in no significant change in Michaelis-Menten parameters *V*_*max*_, *K*_*m,app*_, or *n* (data not shown). These results indicate that we can confidently assign the fitted kinetic parameters to a *rrnA*P3-derived product under these experimental conditions.

### Steady-state measurements show expected dependence on both RNAP and DNA concentration

In bacterial cells there exists a large molar excess of genomic DNA relative to the amount of free RNA, making the *in vivo* global rate of transcription largely independent of gene concentration (49–52). Under these conditions, if the free RNAP concentration is well below the *K*_*m*_, the initial rate becomes independent of DNA concentration and proportional to the free RNAP concentration. As a result, the specific ratios of total RNAP concentration, free RNAP concentration (i.e., RNAPs that are not non-specifically bound to DNA or actively performing transcription (53, 54)) and the number of possible interaction sites on the DNA will determine the reaction rate. To illustrate these concepts *in vitro*, we performed DNA titrations at multiple RNAP concentrations, using both plasmid (2551 bp) and linear templates (250 bp) to provide different numbers of non-specific interaction sites for the RNAP.

Previous measurements using an aptamer-based assay with T7 RNAP illustrated that the DNA concentration dependencies of steady-state rates followed a hyperbolic curve and could be described by Michaelis-Menten kinetics (36). We performed titrations of the *rrnA*P3 plasmid (0.1 – 50 nM) at two different concentrations of *Mtb* RNAP (20 and 100 nM) (**Figure 4A**,**B**) and observed significantly higher rates at the higher RNAP concentration (**Figure 4C**) as expected given the excess of RNAP over DNA. However, we did not observe a monotonic increase in the rates as a function of DNA concentration at either RNAP concentration. At low nM concentrations of the plasmid, we observed the expected increase in steady-state rate; however, as the plasmid concentration was increased, the rate exhibited a concentration-dependent decrease rather than a plateau (**Figure 4C**). Of note, the peak velocity at both concentrations of RNAP occurred at a similar DNA:RNAP ratio, roughly when RNAP was in 10-fold excess of plasmid DNA (**Figure 4D**). These observations are consistent with a mechanism where promoter-specific DNA binding dominates in conditions of limiting DNA concentration, but as the concentration of non-specific binding sites is increased, RNAP binds to sites other than the promoter more often, resulting in a reduction in activity.

**Figure 4:**
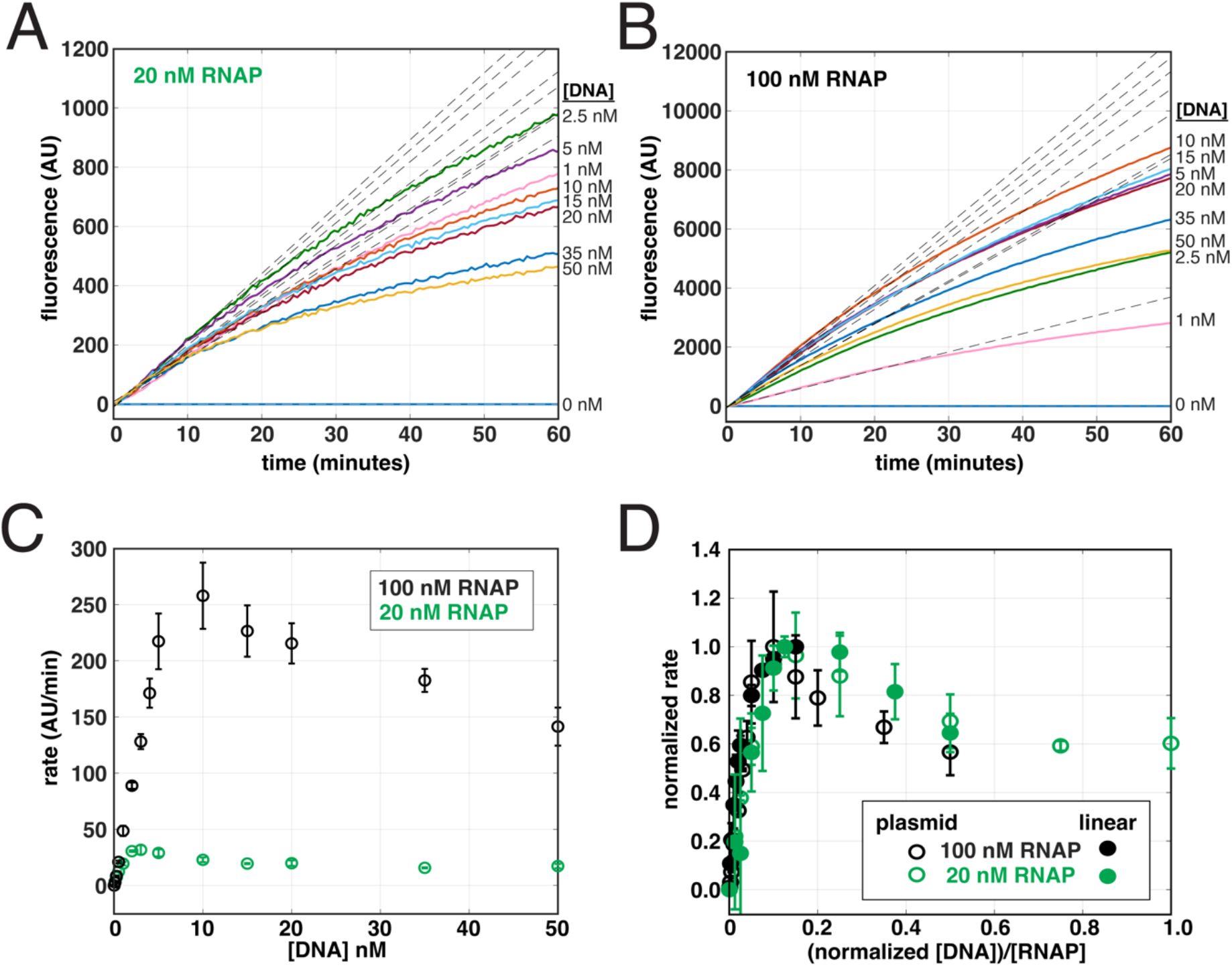
Steady-state rates exhibit a biphasic DNA-concentration-dependence at multiple RNAP concentrations. Real-time data obtained at 500 μM all NTPs, titrating *Mtb rrnA*P3 plasmid DNA template (0.1 – 50 nM) at either **(A)** 20 nM or **(B)** 100 nM *Mtb* RNAP concentrations. The unbiased linear fits of the early times are shown in grey dotted lines for each trace. **(C)** Steady**-**state rates obtained from the linear fits in (A) and (B) for 20 nM (green) and 100 nM (black) RNAP, plotted as a function of *rrnA*P3 plasmid DNA concentration. **(D)** Steady-state rates, normalized from zero to one based on the lowest and highest rate obtained at each RNAP concentration, plotted as a function of the normalized [DNA]:[RNAP] concentration ratios. Included titrations of steady-state rate data are those obtained from the plasmid data (open circles) and from linear data (**Supplementary Figure 6**, closed circles). Linear DNA concentration was normalized to that of the plasmid by dividing by a factor of ten to account for the template length (linear template = 250 bp; plasmid template = 2551 bp).

Analogous titrations done on linear templates containing the *rrnA*P3 promoter sequence (see Methods and **Supplementary Table 2** for template preparation, description, and sequence), (**Supplementary Figure 6A**,**B**) displayed a less prevalent reduction in steady-state rates at higher DNA concentrations (**Supplementary Figure 6C**,**D**). Specifically, the reduction in steady-state rate did not occur until reaching a DNA:RNAP ratio of ∼2:1 for the 20 nM RNAP condition (**Supplementary Figure 6D**) and no reduction was observed in the data collected with 100 nM RNAP over the DNA concentration range tested (**Supplementary Figure 6C**,**D**). On linear templates, the maximal transcriptional activity occurred at a lower excess of RNAP (i.e., higher DNA:RNAP ratio) compared to plasmid templates (**Supplementary Figure 6D**). A shift to a lower DNA:RNAP ratio of the peak velocity may be caused by an increase in non-specific sites. When we normalize the DNA concentration by length, the maximal activities for both linear and plasmid templates overlayed, suggesting that the rate decrease is due to non-specific binding (**Figure 4D**). Combined, these experiments demonstrate quantification of the dependencies of the steady-state rates on both RNAP and DNA concentration, as well as highlight the advantages of using low DNA concentrations when quantifying promoter-specific initiation rates. This is especially important on plasmid templates which typically contain a higher number of possible non-specific interaction sites.

### Quantitative agreement between fluorescence and gel-based approaches

Our fluorescence measurements clearly exhibit a promoter dependence (**Figure 3B**). As expected, performing analogous low throughput experiments involving incorporation of ^32^P labeled UTP followed by separation via polyacrylamide gel electrophoresis (Methods) revealed a specific band of the expected length in accordance with the position of the termination sequence on the plasmid containing the *rrnA*P3 promoter but not on the promoterLESS plasmid template (**Figure 5A**). In addition, bands larger than the full-length product were observed with both templates (**Supplementary Figure S7A**). As these bands were of equal intensity in both the *rrnA*P3 and promoterLESS data, we concluded that their generation was due to a component of the plasmid other than the promoter region being studied and they are not considered further.

**Figure 5:**
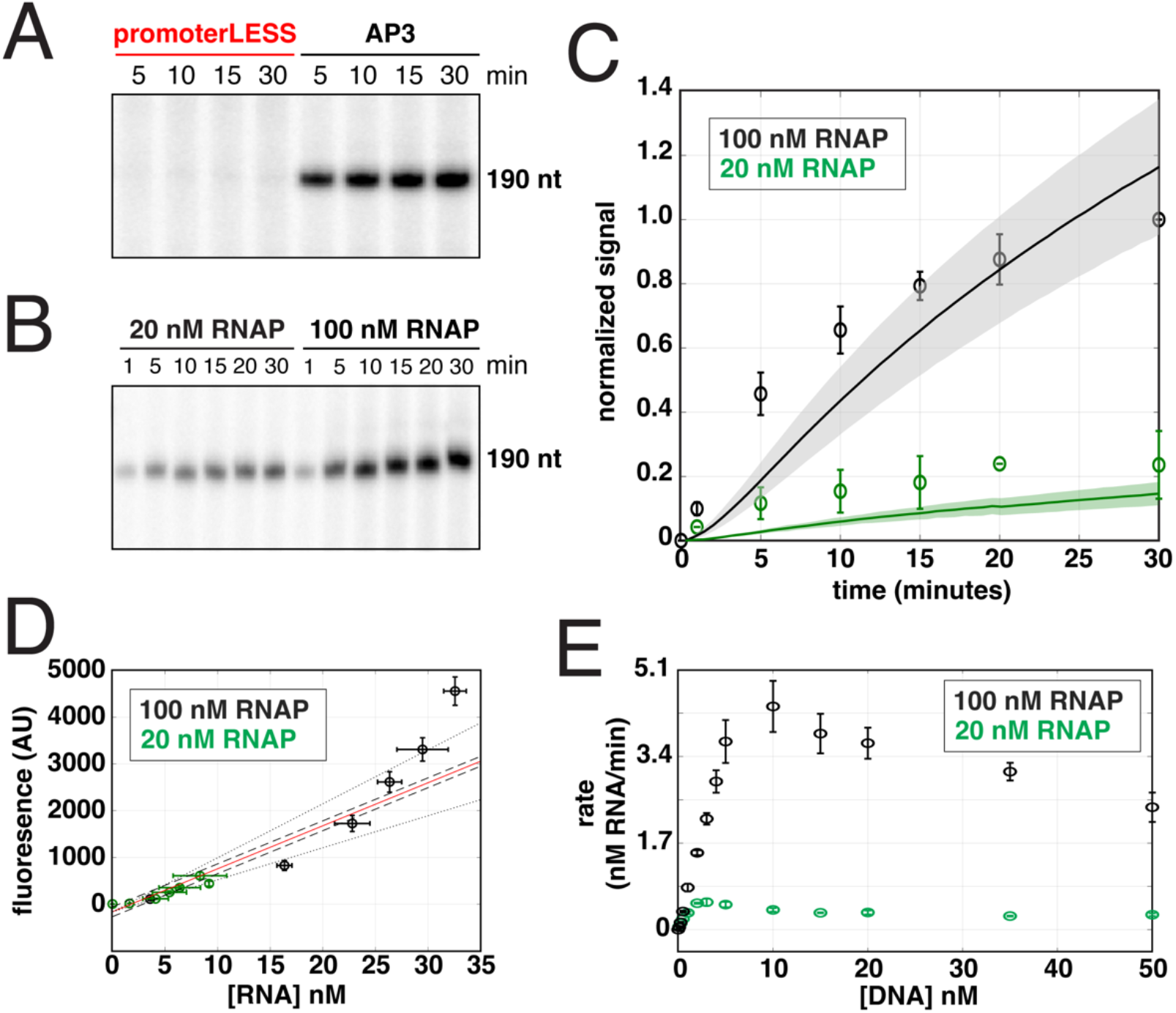
Gel-based experiments illustrate quantitative agreement with fluorescent-based kinetics. Radioactive transcription gels showing the time evolution of the full-length transcript band (see **Supplementary Figure 7** for full gel images) obtained with **(A)** 100 nM *Mtb* RNAP with 5 nM of either the promoterLESS or *rrnA*P3 plasmid templates and **(B)** 20 nM and 100 nM *Mtb* RNAP with 5 nM *rrnA*P3 plasmid template. **(C)** Comparison of time-dependent signals from the fluorescent-aptamer-based assay (lines) and the gel assay (open circles) as a function of time for the 20 nM and 100 nM RNAP. The gel data was normalized by the 30 minute signal at 100 nM RNAP concentration. The fluorescent data was normalized by the average ratio of the fluorescent and gel data at the 15, 20, and 30 minute time points. **(D)** Fluorescent signal plotted against RNA concentration from the data shown in (B). A linear fit (red) along with 95% confidence regions for both the intercept (dashed lines) and the slope (dotted lines) were used to estimate a calibration factor. **(E)** Data from **Figure 4C** replotted using the calibration factor from (D) to convert the y-axis from arbitrary fluorescence units per minute to [RNA]/minute. Error bars indicate the standard deviations from three – four biological replicates for fluorescence data and two – three biological replicates for gel-based data in all sub-plots.

We next asked whether the time evolution of the promoter-specific band yielded the same kinetics as those measured in real-time via the aptamer assay. For this comparison, gel-based experiments were performed at 20 and 100 nM *Mtb* RNAP with 5 nM *rrnA*P3 plasmid DNA (**Figure 5B**; **Supplementary Figure S7B**). Comparing the relative rates across the two different RNAP concentrations showed roughly the same RNAP concentration dependence for both the fluorescence and ^32^P incorporation approaches (**Figure 5C**). This agreement provides further evidence that the fluorescence signal reports on the promoter-specific activity of the system. Furthermore, it suggests that one may use the calculated RNA concentration of the specific, promoter-derived band from gel quantifications to convert the arbitrary fluorescence signal into an absolute RNA concentration. After constructing a calibration curve linking fluorescence and RNA concentration (**Figure 5D**), we converted the raw data and the steady-state rate measurements into [RNA]/time **(Figure 5E)**, providing meaningful units to the *V*_*max*_ value. We note this conversion factor will not be universal and will depend highly on the experimental setup and conditions (see **Supplementary Discussion**). In summary, results from the gel-based and fluorescence-based assays are consistent, and performing both assays provides a means of converting the arbitrary fluorescence signal into absolute RNA concentration.

### The spinach aptamer and DFHBI have no effect on the measured steady-state rates

We have demonstrated that use of the aptamer assay allows for the quantification of steady-state rates of transcription. However, the two additional elements that could theoretically alter the measured rate have not been discussed: the aptamer sequence and the DFHBI dye. To examine the effect of the aptamer sequence or the presence of the dye on the rate of transcription, we performed ^32^P incorporation assay measurements on plasmid templates containing the *rrnA*P3 promoter in the presence and absence of the aptamer sequence and in the presence and absence of dye. We observed roughly the same steady-state rate ± aptamer sequence (**Supplementary Figure 8A**,**B**), suggesting that the fluorescence assay can be taken to report on the rates of transcript production without effects from the aptamer. Additionally, in titrating DFHBI concentration up to 20 µM (the concentration used in the fluorescence assay) in ^32^P incorporation experiments, we observed no change in the amount of RNA generated 30 min after initiating the transcription reaction (**Supplementary Figure 8C**,**D**), consistent with previous measurements made with T7 RNAP (36).

We note that in aptamer assay, we observe a change in the magnitude of the fluorescent signal when titrating dye concentrations in the context of the same transcription reaction (**Supplementary Figure 9A**). This can be explained by the finite affinity of the dye for the aptamer. As dye concentration is increased through the K_d_, higher and higher fractions of aptamer are bound, effectively increasing the gain of the signal. However, if experiments are always performed at a fixed dye concentration, preferably higher than the binding affinity to maximize signal (**Supplementary Figure 9B**), comparing fold-changes in fluorescence is a valid approach under steady-state conditions (**Supplementary Figure 9C**,**D**).

### High-throughput capabilities of the fluorescent assay permits concentration dependencies of individual NTPs to be measured in a single experiment

Higher concentrations of the initiating nucleotide (iNTP) than the subsequent NTPs are frequently required for promoter specific initiation, especially on ribosomal RNA promoters where incorporation of the iNTP increases the population of open complexes at equilibrium, which can be a rate-limiting step (17, 55). We hypothesized that when measured under steady-state conditions, titrations of the iNTP would thus yield the highest *K*_*m*_ compared to the other NTPs on the *Mtb* ribosomal RNA promoter. To measure the dependence of steady-state rates on the concentration of individual NTPs on the *rrnA*P3 plasmid template, we performed titrations of each NTP from 2.5 – 500 µM in a background of saturating concentrations (500 µM) of the other three NTPs (**Figure 6A**). Each biological replicate incorporated three technical replicates across ten concentrations per individual NTP titration resulting in a total of 120 total kinetic measurements obtained in parallel, highlighting the high-throughput nature of this assay. The resulting traces were fit to extract steady-state rates which were fit as a function of individual NTP concentration (**Figure 6B**). In cases where the identity of either the first or the first two nucleotides transcribed were in limiting conditions (i.e., the titration GTP or the titration of all NTPs together), a simple Michaelis-Menten equation (**Eq. 1**) did not generate good fits. For these titrations **Eq. 2** was used to account for the sigmoidal dependence. The *V*_*max*_ for each titration was within error, as at the high concentrations of the titrated NTP, all conditions become identical (**Figure 6A,B**; **Supplementary Table 3**). As predicted, the data revealed that the transcription rate depends most strongly on the iNTP (GTP) concentration with a *K*_*m,app*_ of 16 ± 2 µM, consistent with the results from single time-point experiments on *E. coli* ribosomal RNA promoters (55). As the identity of titrated nucleotide moved further downstream within the sequence of the initially transcribed region of the promoter, the measured *K*_*m,app*_ shifted to lower concentrations (**Figure 6B**; **Supplementary Table 3**). Notably, these trends in *K*_*m,app*_ and *V*_*max*_ as a function of nucleotide identity were recapitulated using linear templates containing the *rrnA*P3 sequence (**Figure 6C,D**) although the *K*_*m,app*_ values were higher in all cases (**Figure 6D**; **Supplementary Table 3**), consistent with previous linear and plasmid template comparisons made on ribosomal RNA promoters (55, 56). Even though we observed a substantially lower *V*_*max*_ when considering the scale of arbitrary fluorescent units on linear as opposed to plasmid templates, the fact these nucleotide-identity-dependent trends could be easily measured on either template, including the sigmoidal concentration dependencies, demonstrates the utility of the assay regardless of the exact experimental setup.

**Figure 6:**
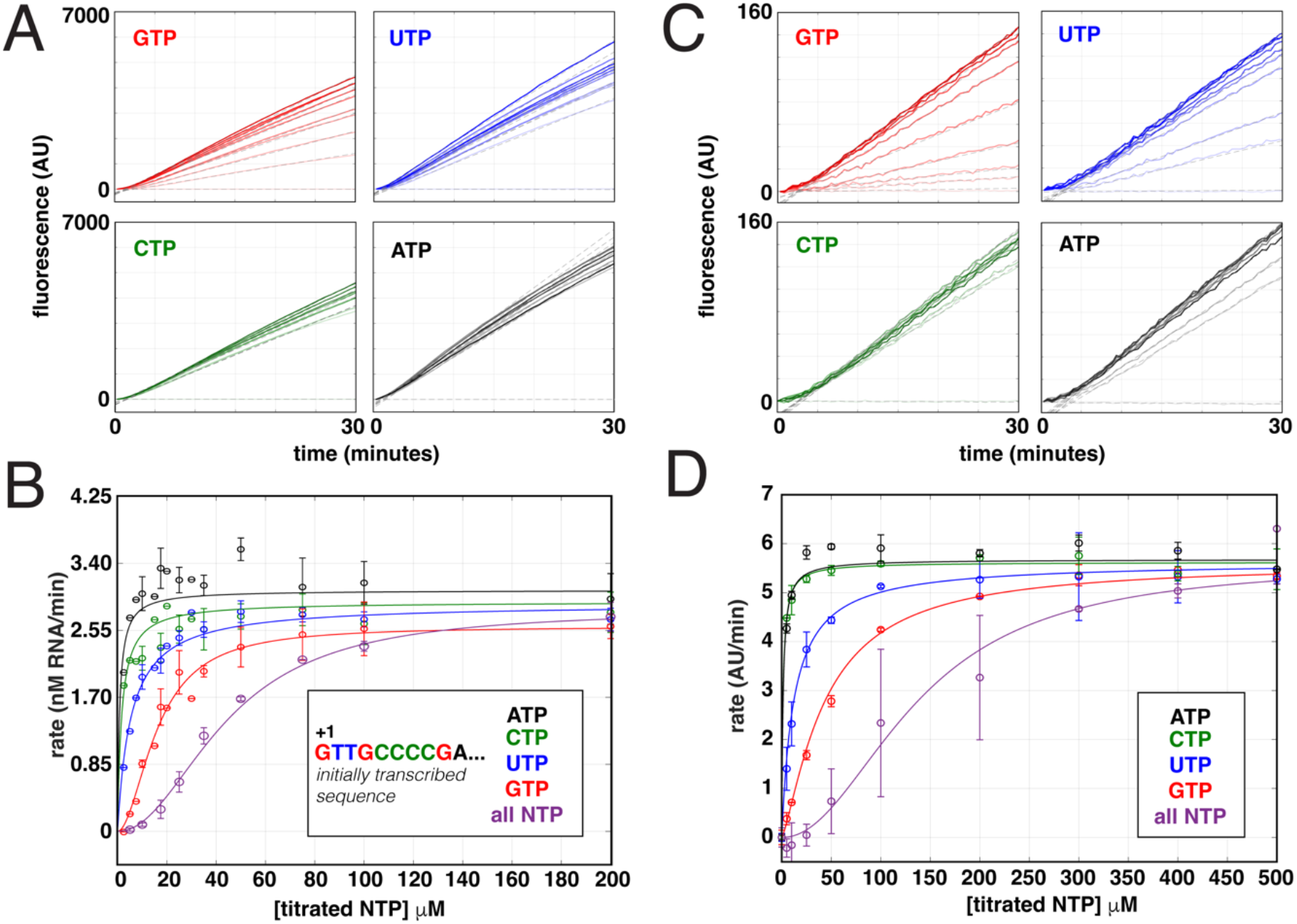
Titrations of individual NTPs reveal incorporation of the initiating nucleotide is rate limiting. **(A)** Real-time fluorescent traces and linear fits obtained using 100 nM *Mtb* RNAP, 5 nM *rrnA*P3 plasmid DNA, and individually titrating GTP, UTP, CTP, or ATP in the presence of 500 μM of the other three, non-titrated NTPs. **(B)** Rate dependence on the concentration of individual NTPs from the data in (A) as well as the titration of all NTPs equally. Note that for clarity, only the titration data out to 200 μM NTPs is shown. The y-axis was converted to report on absolute RNA concentrations using a conversion factor of 58.8 fluorescence units/ 1 nM RNA, based on the calibration in **Figure 5D. (C)** Real-time fluorescent traces and fits as in (A), except using 25 nM of the linear *rrnA*P3 DNA template. **(D)** Rate dependence on the concentration of individual NTPs from the data in (C) as well as the titration of all NTPs equally. Error bars in (C) and (D) represent standard deviations of two – four biological replicates each with two – three technical replicates. Rate dependencies in (C) and (D) were fit to **Eq. 1** or **Eq. 2** depending on the identity of the titrated NTP, and fitted parameters are summarized in **Supplementary Table 3**.

### Quantification of transcription factor activity under steady-state conditions

When evaluating the effect of a transcription factor *in vitro*, the most common approach is to use single timepoint measurements to compare transcription in the presence and absence of the factor. However, by only evaluating a single timepoint for each condition there is no guarantee or confirmation that the measurements are representative of changes occurring throughout a steady-state process. In contrast, the aptamer assay directly measures the effect of transcription factors in real-time and by focusing the analysis on the linear regime in an unbiased manner, ensures that factor-dependent changes are quantified under steady-state conditions. To demonstrate the use of the assay in analysis of transcription factor effects, we turned to two well-studied *Mtb* transcription factors, CarD and RbpA.

Previous studies have shown that both CarD and RbpA stabilize the open complex (15, 16, 57, 58), slow promoter escape (17) and activate transcription from the *rrnA*P3 promoter (57, 59). In addition, these factors are known to bind the initiation complex cooperatively and act synergistically (16, 17, 58). We measured transcription in the presence of CarD, RbpA, or both at saturating concentrations on the linear *rrnA*P3 template. We monitored fluorescence over time and fit the traces to obtain steady-state rates (**Figure 7A**). Between two – five biological replicates were used to calculate and compare the average rates at each condition. Consistent with previous work, we observed a 2.8-fold, 4-fold, and 9.9-fold increase in the rate of RNA production in the presence of RbpA, CarD, and both factors together, respectively (**Figure 7B**).

**Figure 7:**
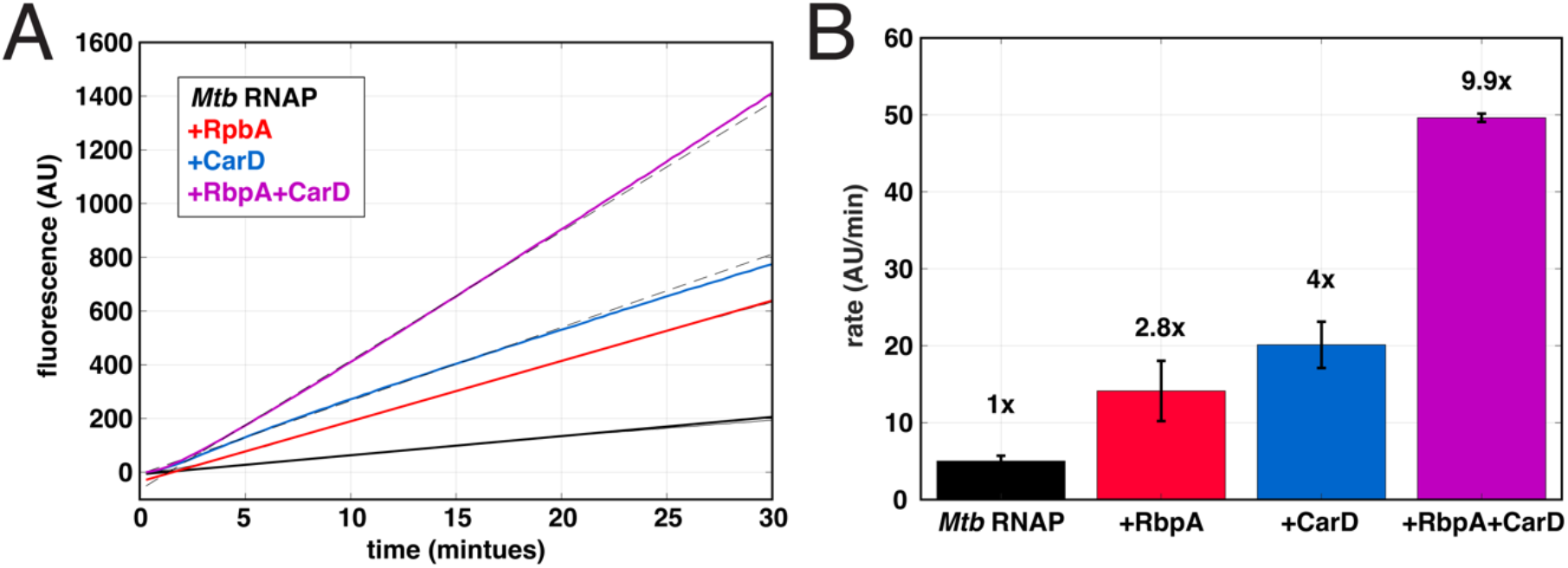
Factor-dependent effects on steady-state transcription rates. **(A)** Real-time fluorescent traces collected with 100 nM *Mtb* RNAP, 500 μM all NTPs and 25 nM linear *rrnA*P3 template in the presence of saturating *Mtb* CarD and RbpA (solid curves) with the associated linear fits (dotted lines). **(B)** Comparison of steady-state rates (average of two – five biological replicates) reveals the extent of transcriptional activation by RbpA (red), CarD (blue), and the combination of RbpA and CarD (purple). Fold changes in the steady-state rate relative to *Mtb* RNAP alone are indicated above each condition.

### Quantification of inhibitory concentrations of antibiotics

Bacterial RNAPs are the direct targets of many antibiotics (reviewed in (8, 60)) and the search for new antibiotics to overcome drug resistance is never-ending. This is particularly important in the battle against *Mtb*, the causative agent of tuberculosis, as multi-drug resistant strains are becoming more prevalent (61). To this end, the development of new small molecule inhibitors that work at the level of transcription is of high interest. Here, we demonstrate that the aptamer assay can be used to quantify antibiotic-dependent inhibition of steady-state rates. We illustrate this with the well-characterized antibiotics Rifampicin and Fidaxomicin, currently used to treat *Mtb* and *Clostridium difficile* infections by directly targeting the bacterial RNAP (62–65).

Titrations of both Rifampicin and Fidaxomicin were performed using either 100 nM *Mtb* or *E. coli* RNAP with 5 nM *Mtb rrnA*P3 plasmid template (Methods), and the kinetic traces were fit to extract the steady-state rates. Although *E. coli* RNAP exhibits a lower maximal rate of transcription on this template, as inferred from the fluorescence signal in the absence of antibiotic (**Supplementary Figure S10**), for both the RNAPs, we observed a concentration-dependent decrease in the measured steady-state rates upon increasing antibiotic concentrations (**Figure 8**). Fitting the data to an inhibition curve (**Eq. 3**) permitted the calculation of half-maximal inhibitory concentrations. Consistent with previous work, the *IC*_*50*_ for Rifampicin on *E. coli* RNAP (15 ± 2 nM) and *Mtb* RNAP (17 ± 2 nM) were within error (**Figure 8A**) (57, 66). In addition, the same analysis for Fidaxomicin yielded an approximately three-fold higher *IC*_*50*_ on *E. coli* RNAP (400 ± 110 nM) than on *Mtb* RNAP (138 ± 92 nM) (**Figure 8B**), consistent with previous reports of *Mtb* being more sensitive to the drug than *E. coli* (63, 67).

**Figure 8:**
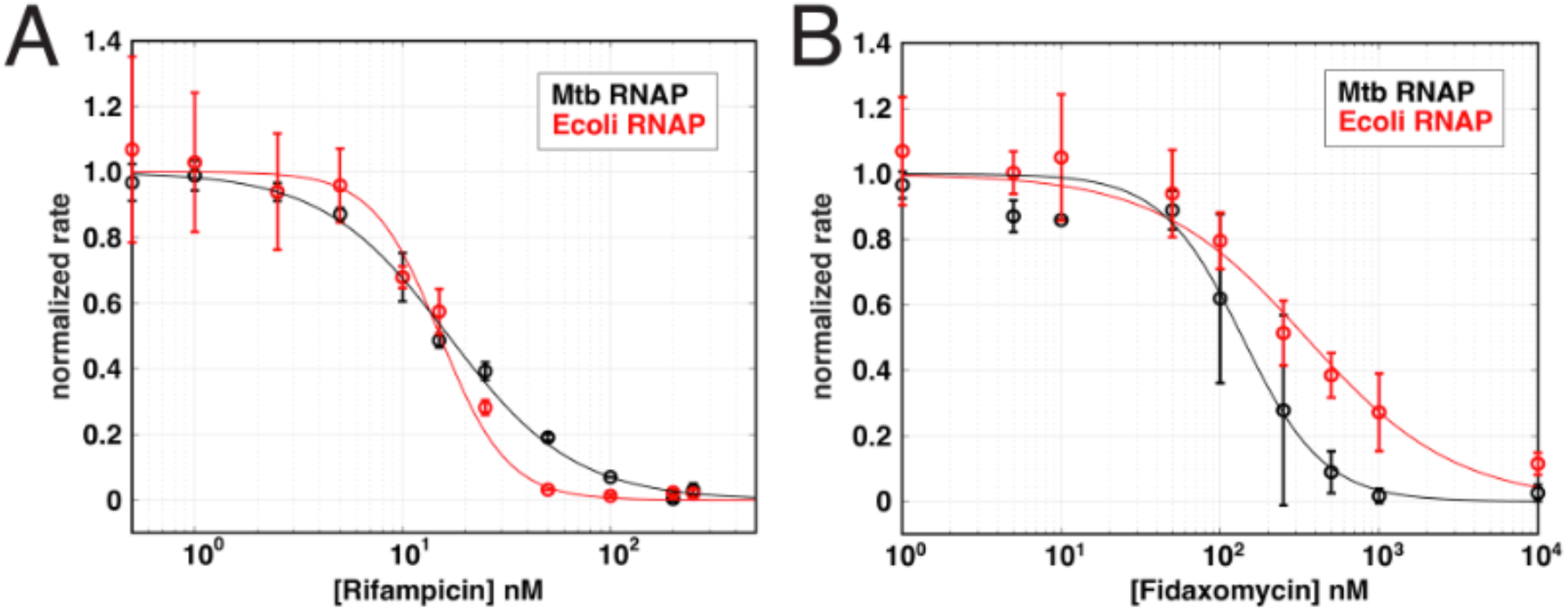
Quantification of antibiotic IC_50_ values based on changes in observed steady-state rates. Normalized steady-state rates plotted as a function of **(A)** Rifampicin and **(B)** Fidaxomicin concentrations for *Mtb* (black) and *E. coli* (red) RNAP. All experiments were performed using 1 mM of each NTP and 5 nM *rrnA*P3 plasmid DNA. See **Supplementary Figure 10** for the corresponding real-time data, linear fits, and un-normalized rates. The normalized steady-state rates are based on the associated fits using **Eq. 3** (**Supplementary Figure 10E**,**F**).

## Discussion

### High-throughput kinetic measurements of full-length transcription

The fluorescence experiments presented here are an attractive alternative technique to the standard radiolabeled-nucleotide incorporated gel-based approaches, especially when used in a plate-reader format as described. Hundreds of conditions can be measured in real-time with several technical replicates in a single experiment. We estimate that the throughput is on the order of a hundred times that of standard gel-based approaches, where the real-time measurements facilitate a more precise and accurate determination and quantification of the steady-state regime. As the steady-state rate of full-length transcription (along with the RNA degradation rate) is a crucial metric when considering the flux of biological processes that use RNA as a substrate (1), use of this assay will significantly facilitate the exploration of diverse mechanisms of gene regulation with biological relevance.

We emphasize that radioactive techniques are advantageous when looking to resolve individual RNA transcripts, as well as to determine the relative lengths and amounts of abortive products (68). In fact, the first steady-state assays that reported on bacterial transcription were made under conditions where NTP synthesis was restricted to abortive transcription (69, 70). However, as RNAP does not escape the promoter under these circumstances, these assays did not report on the overall rate of initiation, but rather the rate-limiting step involved in binding and isomerization to the catalytically active open complex (70). Fluorescent-labeled nucleotides that are incorporated into the RNA transcripts have also been used to monitor the steady-state rates of abortive RNA generation with *E. coli* RNAP (71–73), representing an alternative to radioactivity-based approaches. However, this method cannot discriminate between abortive and full-length RNA products. In comparison to other techniques, the fluorescent-aptamer-based assay described here is advantageous, as the observed fluorescence signal reports specifically on full-length RNA products. This is demonstrated in conditions where single NTPs are not included, resulting in the inability of RNAP to escape the promoter and a lack of fluorescence signal change (**Figure 6**). We note that the effects of abortive synthesis are still borne out in the kinetics, as the process of promoter escape can be rate-limiting on some promoters and affected by the initially transcribed sequence (74, 75), which we have included in our DNA constructs. As a result, the assay permits a straightforward way to measure the rate of full-length transcript production dictated by the overall rate of initiation.

We have illustrated that the assay can easily quantitate difference in kinetics due to promoter sequence (**Figure 3**), NTP concentration and identity (**Figure 6**), and RNAP and promoter DNA concentrations (**Figure 4**; **Supplementary Figure 6**). The non-monotonic rate-dependencies observed when titrating DNA (**Figure 4**; **Supplementary Figure 6**) suggest one could use this assay to evaluate how the presence of other promoters or number of non-specific sites compete for RNAP binding and affect the steady-states rates from a particular promoter of interest. Furthermore, in addition to having great utility in multi-round assays, the assay can be used in single-round conditions to evaluate rates of initiation processes such as promoter escape (**Supplementary Figure 3**). The single-round approach has previously been well-described and used for monitoring co-transcriptional RNA folding processes (40, 43). We stress that meaningful quantifications of relative changes between conditions can be made using arbitrary fluorescence units. However, we have also shown the approach and necessary controls to calibrate the fluorescent signal using an independent measure of RNA concentration from identical reactions (**Figure 5**). Further thoughts on calibration and interpretation of the fluorescent signal can be found in the **Supplementary Discussion**.

### Nucleotide-dependent kinetics

By performing titrations of NTP substrates, we have illustrated that *K*_*m,app*_ and *V*_*max*_ parameters can be obtained with a high level of precision under steady-state conditions (**Figure 6**). Using these data, we observed that not all titrations could be fit to a hyperbolic Michaelis-Menten model. Rather, sigmoidal trends were observed for both plasmid and linear DNA templates in the titration of all NTPs together and GTP in the background of saturating concentrations of ATP, UTP, and CTP (**Figure 6B,D**; **Supplementary Table 3**). Initiation requires the binding of the first two nucleotides to form the first RNA phosphodiester bond and, depending on the conformation of the open DNA, base-stacking with the +1/+2 DNA sequences on the template strand may represent a rate-limiting step (14). Our results are consistent with this hypothesis in that the binding of the concentration of the first two NTP substrates has a larger effect on transcription from *rrnA*P3 than the remaining two NTPs (**Figure 6B,D**; **Supplementary Table 3**). For monomeric enzymes containing a single active site, a sigmoidal dependence of velocity on substrate concentration has been referred to as kinetic cooperativity and was first measured with glucokinase (76, 77). Additionally, other mechanisms have been suggested to lead to kinetic cooperativity without direct interactions between binding sites (reviewed in (78)), including the existence of a slow conformational change that precedes substrate incorporation, the existence of multiple enzyme classes capable of binding initiating nucleotides with different affinities, and/or a substrate-induced conformational change to the catalytically active enzyme (78, 79). These models may not be mutually exclusive and, in fact, are supported by numerous experimental studies evaluating the initial nucleotide incorporation process with *E. coli* RNAP (9, 10, 13, 14). While we cannot directly comment on the specific mechanism(s) that results in this apparent positive cooperativity with *Mtb* RNAP, the detection of this sigmoidal behavior would not have been possible without the fine sampling of NTP concentrations and precise measurements of steady-state rates facilitated by the ease and throughput of the aptamer-based assay.

### Future uses for high-throughput detection of basal and regulated transcription kinetics

Given these fluorescent experiments were all performed in a 384 well plate, the assay lends itself naturally to questions that require large data sets. We conclude by summarizing a few exciting possible directions.

Using sequence to predict transcriptional activity remains a challenge, as the context of the entire promoter must be considered when examining how sequence affects initiation kinetics (3). Using the aptamer-based assay, one could generate promoter libraries where sequence context is investigated by introducing sequence mutations in promoter regions of interest (41, 74, 80). Alternatively, one could design and measure basal and regulated transcription rates on large numbers of genomic promoter sequences. This would allow for a side-by-side comparison with genome-wide, *in vivo* measurements that sample the steady-state amounts (i.e., RNA-seq) or the rates of production (i.e., GRO-seq) of transcripts over the entire genome. By comparing *in vitro* regulation kinetics to *in vivo* sequencing data, one could parse out regulation due to the direct effects of a particular transcription factor or small molecule effector from those that are secondary effects generated from *in vivo* cellular perturbations. A tour-de-force example along these lines performed with single-time point, radioactive gel-based approaches can be seen in the study of ppGpp regulation in *E. coli*. Here, 134 promoters were assessed *in vitro*, and the resultant factor-dependent fold change was compared to the *in vivo* response of bacterial strains which were specifically designed to assess the transcriptomic effects of ppGpp directly binding to the RNAP (81). Use of the aptamer-based assay will greatly facilitate the ability to quantitatively probe the kinetics of larger genomic promoter libraries *in vitro* to expand our knowledge of gene regulatory mechanisms.

As we illustrated the utility of the fluorescence assay in measuring antibiotic inhibition (**Figure 8**; **Supplementary Figure 10**), the high-throughput screening of novel inhibitors against the bacterial transcription machinery may also benefit from this assay. Inhibitors that act at the level of initiation could be identified by a reduction the initial steady-state rate, as observed for Rifampicin and Fidaxomicin (**Figure 8**; **Supplementary Figure 10**). In addition, inhibitors of elongation or termination that act to reduce the active RNAP concentration or the free promoter concentration may lead to faster reductions in rate at longer-timescales. As a result, use of this assay under multi-round conditions may permit the identification of compounds that inhibit transcription via different mechanisms. Furthermore, because one tracks the time-dependent changes in fluorescence and not single-time points, compounds that exhibit fluorescent spectral properties are less likely to produce artifactual results. Compounds identified as potential hits can then easily be analyzed via titrations to measure the IC_50_s as described here. Furthermore, to develop structural hypotheses for the mechanism of transcription initiation inhibitors (or classes of inhibitors), titrations of NTPs or promoter DNA could be performed to determine the type of inhibition (e.g., competitive, non-competitive, or uncompetitive) via quantification of changes in *K*_*m,app*_ and/or *V*_*max*_ (82).

## Materials and Methods

### Preparation of DNA constructs

#### Plasmid constructs

Plasmids constructs were ordered from Twist Bioscience (San Francisco, CA), where the sequences of interest were inserted into to the pTwist, High Copy, Amp^R^ plasmid. Plasmids were isolated using a QIAGEN Plasmid Midi Kit (Qiagen; catalog #12123) and concentrations were calculated using the known molecular weights based on each unique insert sequence. The insert sequence corresponding to the *Mtb* ribosomal RNA promoter (*rrnA*P3) contains both the Spinach-mini aptamer sequence (30) and the *E. coli rrnB*P1 *T*_1_ terminator sequence (83) (illustrated in **Figure 1**). The *Mtb rrnA*P3 genomic sequence is present both upstream (from – 60 to +31) and downstream (from +32 to +70) of the aptamer. For additional sequences and further descriptions, see **Supplementary Table 1**.

#### Linear constructs

Linear templates of 250 base-pair (bp) in length, containing the *Mtb rrnA*P3 and Spinach-mini aptamer sequences, were also used for steady-state rate measurements. Two, 200 nucleotide (nt) oligonucleotides were ordered from Integrated DNA Technologies, Inc. (Coralville, IA) as PAGE purified DNA Ultramers™. These oligos contained 150 bp of overlapping sequence, leaving 50 nt overhangs on each end. Annealing and end-filling to generate a 250 bp construct was accomplished with Platinum Taq DNA Polymerase (Invitrogen; catalog # 15966005). These templates underwent an additional PCR step with Klentaq LA DNA Polymerase (84) (DNA Polymerase Technology; catalog # 110), using a biotinylated primer (Integrated DNA Technologies, Inc.). Following each PCR step, samples were purified with QIAquick PCR Purification Kit (Qiagen; catalog # 28104) and purity was verified by Native-PAGE. The final product contained a biotin molecule on the upstream end of the template with the same orientation of *rrnA*P3 and the aptamer sequences as used in the plasmid constructs. Concentrations were calculated using the known molecular weight based on sequence. For primer and final construct sequences, as well as further preparation details, see **Supplementary Table 2**.

#### Protein expression and purification

Plasmids for the co-expression and purification of the *Mtb* RNAP σ^A^ holoenzyme were obtained from Jayanta Mukhopadhyay (Bose Institute) (85), using previously described methods for expression (86). Purification of the holoenzyme complex was performed with methods previously used for *Mycobacterium bovis* core RNAP (15). *Mtb* CarD and RbpA, in pET-SUMO plasmid vectors, were expressed and purified, and the His-SUMO tag was subsequently cleaved in accordance with methods previously described (15, 86). *E. coli* RNAP σ^70^ holoenzyme was purchased from New England BioLabs, Inc. at 1.7 μM concentration (Ipswich, MA; catalog # M0551S). Purified *E. coli* GreB was a generous gift from Irina Artsimovitch (The Ohio State University) (87). Protein concentrations were determined via absorbance using the following extinction coefficients at 280 nm: *Mtb* RNAP σ^A^ holoenzyme (280,452 M^-1^cm^-1^), *Mtb* CarD (16,900 M^-1^cm^-1^), *Mtb* RbpA (13,980 M^-1^cm^-1^), and *E. coli* GreB (36,900 M^-1^cm^-1^).

#### Reagents for transcription assays

All transcription reactions were performed with NTPs (Thermo Scientific; catalog #s: ATP – R0441, CTP – R0451, GTP – R0461, and UTP – R0471), 3,5-difluoro-4-hydroxybenzylidene imidazolinone (DFHBI) dye (Sigma Aldrich; catalog # SML1627), and RiboLock RNase inhibitors (Thermo Scientific; catalog # EO0381). DFHBI concentration, stored in 100 mM HEPES, pH 7.6, was determined by absorbance using an extinction coefficient at 420 nm of 31,611 M^-1^cm^-1^. Salmon-sperm DNA (Invitrogen; catalog # 15632011) was used as a competitor for single-round experiments (17). Rifampicin (Sigma-Aldrich; catalog # R3501) and Fidaxomicin (Selleckchem; catalog # S4227) solids were dissolved in DMSO. The molar concentration of Fidaxomicin was determined by weight and of Rifampicin by absorbance using an extinction coefficient at 470 nm of 15,300 M^-1^cm^-1^ (88).

#### Plate-reader fluorescence measurements

To measure multi-round kinetics under steady-state conditions in real-time, we monitored the change in DFHBI fluorescence upon binding a transcribed, full-length RNA sequence containing the Spinach-mini aptamer. Data was collected using a Synergy™2 multi-detection microplate reader (BioTek Instruments, Inc., Winooski, VT) with the corresponding Gen5™ analysis software. DFHBI fluorescence was measured with a tungsten light source equipped with a 505 nm longpass dichroic mirror, excited with a 460 ± 40 nm bandpass filter, and the resulting emission signal was monitored with a 528 ± 20 nm bandpass filter. Data was acquired at a read height of 7.00 mm, typically in 20 – 30 sec intervals, with varying total acquisitions times, not exceeding 75 min.

Transcription reaction master mixes containing 90% of the final volume included the RNAP holoenzyme, NTPs, DFHBI, and RNase inhibitors. Based on the volumes added for each corresponding buffer addition and concentrated stock component, the final solution conditions are 20 mM Tris (pH 8.0 at 25°C), 40 mM NaCl, 75 mM K-glutamate, 10 mM MgCl_2_, 5 μM ZnCl_2_, 20 μM EDTA, 5% (v/v) glycerol (defined as transcription buffer) with 1 mM DTT and 0.1 mg/mL BSA. From these master mixes, a volume of 9 μL for each technical replicate was transferred to an individual well in a 384 well, low volume, round-bottom, non-binding polystyrene assay plate (Corning; catalog # 4514). For the negative controls, which represent the entirety of the reaction components except for DNA, 1 μL of transcription buffer was added to account for the remaining 10% of the final reaction volume (10 μL). Wells were covered with an optical adhesive to prevent sample evaporation (Applied Biosystems; catalog # 4360954). The plate was then incubated for 10 min at 25°C, followed by a 30 sec shake agitation in the microplate reader. An initial reading of the negative controls was used to scale the automatic gain adjustment to the background signal with an arbitrary value of 1000 RFUs and was applied to all the subsequent reads. The adhesive cover was removed, and the transcription reaction was initiated with 1 μL DNA unless otherwise indicated.

Multichannel pipettors were used to reduce the initiation time difference across wells. Once the DNA was added, the plate was agitated for 15 sec before starting the kinetic measurements. Summarized below are the concentrations used for all fluorescent experiments. Unless otherwise stated, the reaction master mixes always contained 20 μM DFHBI and 0.4 U/μL RNase inhibitors.

#### Titration Experiments

NTP titrations: 100 nM *Mtb* σ^A^ RNAP holoenzyme was pre-incubated with NTP concentrations ranging from 2.5 – 500 μM and the reaction was initiated with either 5 nM *Mtb rrnA*P3 or promoterLESS plasmid DNA constructs, or 25 nM *Mtb rrnA*P3 linear DNA constructs. This 25 nM concentration for linear DNA was chosen as it represents the *K*_*m*_ obtained with DNA titrations at this RNAP concentration (**Supplementary Figure 6C**). Titrations with an individual NTP were done in the presence of 500 μM of the other three NTPs. For titrations of all NTPs, the concentrations of each NTP were varied equally.

DNA titrations: 500 μM all NTPs were pre-incubated with either 20 or 100 nM *Mtb* σ^A^ RNAP holoenzyme. The reaction was initiated with concentrations of 0.1 – 50 nM *Mtb rrnA*P3 plasmid DNA or 2.5 – 150 nM *Mtb rrnA*P3 linear DNA constructs.

DFHBI titrations: DFHBI dye (concentrations ranging from 0.005 – 40 μM) was pre-incubated with 100 nM *Mtb* σ^A^ RNAP holoenzyme and 500 μM all NTPs. The reaction initiated with 5 nM *Mtb rrnA*P3 plasmid DNA. For each dye concentration examined, a negative control (no DNA) was collected to correct for any time-dependent changes in the background fluorescent signal. Antibiotic titrations: Rifampicin (concentrations ranging from 0.5 – 250 nM), or Fidaxomicin (concentrations ranging from 0.0001 – 100 μM) was preincubated with 100 nM *Mtb* σ^A^ or *E. coli* σ^70^ RNAP holoenzyme and 1 mM all NTPs. The reaction initiated with 5 nM *Mtb rrnA*P3 plasmid DNA. The final DMSO concentration was kept under 0.2% (v/v) to avoid any concentration-dependent effects of DMSO on RNA folding (89), which would affect the fluorescent signal.

#### Transcription Factor Experiments

Experiments were performed using saturating concentrations of transcription factors, pre-incubated with 100 nM *E. coli* σ^70^ or *Mtb* σ^A^ RNAP holoenzyme and 500 μM all NTPs. Reactions were initiated with 5 nM *Mtb rrnA*P3 plasmid DNA (*E. coli* GreB) or 25 nM *Mtb rrnA*P3 linear DNA (*Mtb* CarD and RbpA). GreB experiments used 1 μM, which binds *E. coli* RNAP core and σ^70^ holoenzyme with ∼ 10 nM affinity (90) and elongation complexes with ∼ 100 nM affinity (91, 92). CarD and RbpA experiments used 1 and 2 μΜ, respectively, as these concentrations lead to saturable open-complex formation kinetics with the *Mtb* RNAP σ^A^ holoenzyme (15, 58).

#### Single-round experiments

To prevent multiple rounds of RNA generation derived from dissociated/terminated RNAPs rebinding the promoter, a competitor DNA was added. We used either salmon-sperm DNA or a plasmid DNA template that contained the *Mtb rrnA*P3 sequence but lacked the aptamer sequence (**Supplementary Table 1**). Pre-formed promoter-bound complexes (100 nM *Mtb* σ^A^ RNAP holoenzyme and 5 nM *Mtb rrnA*P3 plasmid DNA) with 20 μM DFHBI and 0.4 U/μL RNase inhibitors were initiated with either 75 µg/mL of salmon-sperm DNA or 150 nM “without aptamer” plasmid DNA along with all NTPs at various concentrations. These concentrations of competitors were pre-incubated with RNAP and *Mtb rrnA*P3 plasmid DNA containing the aptamer sequence and caused no change in fluorescence upon initiating the reaction with NTPs (**Supplementary Figure 3A**).

#### Radio-labeled NTP Incorporation Transcription Experiments

Transcription experiments were performed with 500 µM each NTP under identical solution conditions and temperature as multi-round fluorescence experiments. *Mtb* σ^A^ RNAP holoenzyme and DFHBI concentrations were varied and always included unless otherwise indicated. 20 nM of α-^32^P UTP was added to label the nascent RNA transcripts via incorporation by RNAP. Reactions were initiated by addition of 5 nM *Mtb rrnA*P3 or promoterLESS plasmid DNA constructs. 5 µL aliquots were removed from the reaction mixture at 1, 5, 10, 15, 20, and 30 minutes and combined with 5 µL of quench solution containing 95% formamide, 0.015 M EDTA, 0.05% (w/v) xylene cyanol, and 0.05% (w/v) bromophenol blue. 5 µL of each quenched timepoint was then loaded onto a 5% denaturing PAGE gel. Gels were run in TBE for two–three hours at 1500 V and then transferred to a phosporimaging cassette. After exposing for 18 hours, phosphorimaging screens were imaged via a Typhoon 9000 phosphorimager. Bands of interest were quantified using ImageQuant software and converted to reaction RNA concentrations as previously described (5).

#### Simulations of steady-state kinetics

Master-equation based simulations were used to explore the effects of aptamer folding and dye binding rates on the observed steady-state fluorescent signal in **Supplementary Figures 1** and **9C,D**. The kinetic model consisted of rate constants describing RNAP-promoter binding (*k*_*on*_) and dissociation (*k*_*off*_), promoter escape (*k*_*escape*_), aptamer folding (*k*_*folding*_), and dye binding (*k*_*bind*_) and dissociation (*k*_*diss*_) and is shown graphically in **Supplementary Figure 1**. Pseudo first order can be assumed for the binding rate constants for RNAP-DNA (*k*_*on*_) and dye-RNA (*k*_*bind*_), when these reactants are in excess. These rate constants are proportional to concentrations of RNAP and dye, respectively. All other rate constants are uni-molecular and concentration independent. The simulation begins with 100% of the population in the unbound polymerase state (R+P). The promoter escape step regenerates free polymerase and promoter and produces an unfolded RNA transcript. Thus, the polymerase/promoter system can generate multiple transcripts over time and the population of unfolded transcript can increase without limit. The accumulation of dye-bound, folded aptamer is what is plotted as a function of time and is taken as a measure of the expected fluorescent signal in the actual assay. At steady state, the concentrations of R+P, RP, and unfolded aptamer are constant while the concentrations of unbound and bound folded aptamer increase linearly in a constant ratio dictated by the dye affinity and concentration.

### Analysis of fluorescent time-courses

#### Extraction of steady-state rates

As the fluorescent signal in the absence of aptamer formation does not significantly change across conditions where DNA, NTPs, or the aptamer sequence were left out of the reaction (**Supplementary Figure 2**), the same negative control could be used to correct all experimental conditions within a titration. This applies only if the time-course measurements are made under the same solution conditions and instrumental detection scaling. However, to minimize experimental and/or instrumental variation, each time a new experiment was performed, a minimum of two – three technical replicates of the negative control (leaving out DNA) were collected and measured concurrently with the experimental data. Prior to data analysis, the experimental traces underwent two subtractions: first, the fluorescent value recorded at the initial time point was subtracted from all time-points, bringing the starting fluorescent value to zero, and second, the fluorescence from the experimental traces was subtracted using the corresponding signal from each time-point of the negative control. Linear regression of the resulting florescent traces was then performed with a custom Matlab fitting program (described further in Results), which can be found on Github (https://github.com/egalburt/aptamer-flux-fitting). Using a statistical weighting from two – three technical replicates per condition and a user inputted R^2^ value to define the goodness of fit (**Supplementary Figure 5**), an unbiased determination of the linear regime can be obtained, where the slope of the fitted line reports on the steady-state rate in units of change in fluorescence/time. In the work presented here, R^2^-thresholds > 0.998 were used.

#### Non-linear regression analyses of concentration dependencies

For analyses of titration data that yielded hyperbolic concentration-dependencies in steady-state rates, a Michaelis-Menten equation was applied, fitting the data to **Eq. 1**

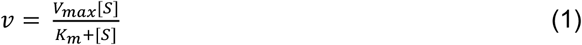

where *v* represents the steady-state rate and *S* represents the titrated substrate, either DNA or individual NTPs. Steady-state rates determined in some NTP titrations were not well-described by **Eq. 1**, displaying sigmoidal concentration-dependences. These titrations were fit to **Eq. 2**, a modified Michaelis-Menten equation with three parameters: *V*_*max*_, an apparent *K*_*m*_ (*K*_*m*,app_), and the exponent *n*.

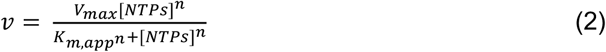

Here, we define *K*_*m*,app_ as the effective half maximal concentration (*EC*_*50*_) of NTPs, which is equivalent to the standard operational definition of *K*_*m*_ when *n* = 1. It should be noted that while this equation is functionally identical to the Hill equation, if a non-unity *n* is the result of the kinetics from a single-site-binding system, then *n* is only effectively demonstrative of the type (positive/negative) and magnitude of cooperativity. In such a case, *n* has no specific physical meaning as it does in more traditional cases such as cooperative binding of multiple ligands (78). The kinetics of transcription inhibition as a function of antibiotic concentration at constant NTP concentrations were also found to be sigmoidal. For determination of the half maximal inhibitory concentration (*IC*_*50*_) of Rifampicin and Fidaxomicin, the concentration-dependent reduction in steady-state rates was fit to **Eq. 3**.

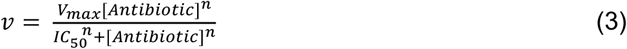

### Statistical Analyses

For all fluorescent data presented unless otherwise indicated, between two – four biological replicates were collected for each condition tested. Each biological replicate contained between two – three technical replicates. The standard deviations from the technical replicates were used as statistical weights during the linear regression analysis used to extract steady-state rates. The rates reported are obtained from the averages and standard deviations of the biological replicates. All non-linear regression analyses were performed using these deviations as a statistical weight. Errors in *V*_*max*_, *K*_*m,app*_, *n*, and *IC*_*50*_ values are those obtained from the fit of the averaged data set. For simplicity, only a single representative biological replicate is shown for the real-time data titration plots. For the gel-based experiments, quantifications of band intensity are calculated from two – three biological replicates and errors are represented as standard deviations.

## Acknowledgments

The authors would like to thank Drs. Dylan Plaskon (University of Wisconsin at Madison) for helpful advice during the infancy of this work, Wayne Barnes (Washington University) for providing advice and reagents for the design of DNA templates, Irina Artsimovitch (The Ohio State University) for providing purified *E. coli* GreB, and Nicole Fazio for critical reading of the manuscript. Additionally, thanks to the Washington University High-Throughput Screening Center (HTSC) staff for technical assistance with plate-reader measurements.

