## Supplemental Information for "High-throughput, fluorescent-aptamer-based measurements of steady-state transcription rates for *Mycobacterium tuberculosis* RNA polymerase"

### This PDF file includes:

**Supplementary Discussion**

Relative vs. absolute RNA concentration measurements: cautions in calibration and interpretation of florescent signal across different experimental conditions

**Supplementary Tables**

**Table S1:** Insert sequences for plasmid DNA constructs

**Table S2:** Primer and final sequences for linear DNA constructs

**Table S3:** Calculated $K_m$, $V_{max}$ and $n$ values from NTP titrations

**Supplementary Figures**

**Figure S1:** Simulations where aptamer-folding is rate-limiting

**Figure S2:** Real-time fluorescent signal in the absence of aptamer formation

**Figure S3:** Single-round trap controls and kinetics

**Figure S4:** GreB affects the long-term behavior of fluorescent traces

**Figure S5:** Overview of variable-time, iterative linear fitting approach

**Figure S6:** Real-time data for DNA titrations on linear templates

**Figure S7:** Full gel images of the data presented in main text Figure 5

**Figure S8:** Gel-based analysis of the effect of the aptamer sequence and DFHBI dye

**Figure S9:** Concentration dependencies of DFHBI on the fluorescent-aptamer signal

**Figure S10:** Real-time data for antibiotic titrations and unnormalized $IC_{50}$ fits

**Supplementary References**

### Supplementary Discussion

#### Relative vs. absolute RNA concentration measurements: cautions in calibration and interpretation of florescent signal across different experimental conditions

Throughout this work, we have illustrated that quantifications of relative changes between conditions can be made using arbitrary fluorescence units. However, we have also shown that the fluorescent signal may be calibrated using an independent measure of RNA concentration from identical reactions. The calibration presented here follows previously described methodologies (1, 2), and uses the signal intensity of the full-length transcript band resolved from gels (**Figure 5B**) to convert RNA concentration using a standard curve (**Figure 5D**). This approach depends on identification of a promoter-specific transcript of the correct length (**Figure 5A**; **Supplementary Figure 7A**). We note that if the total incorporated radioactivity signal were to be used, an overestimate of the promoter-derived, aptamer containing RNA concentration would result, as can be seen by the presence of non-promoter-derived transcripts (**Supplementary Figure 7A**). Similarly, one would also likely overestimate the RNA aptamer concentration if measuring RNA concentration via spectroscopic approaches, such as monitoring absorbance at 260 nm following DNase treatment or monitoring the fluorescence of RNA-specific interacting dyes, as both these approaches would measure the total RNA concentration and not just the promoter-derived product containing the aptamer.

Other studies have bypassed gel-based approaches completely and suggested to calibrate RNA amounts by comparing the fluorescent signal of the experiment to that of known concentrations of purified aptamer (3). While attractive in theory due to its simplicity, this assumes that the equilibrium fraction of folded aptamer when purified is the same when co-transcriptionally folded and generated *in situ*. Here, one must use caution as the specific method used for *in vitro* folding can affect the equilibrium amount of folded aptamer that is capable of binding dye and as a result dictate the fluorescent signal (4). In fact, previous calibration attempts confirmed this assumption, where the calibration based on aptamer fluorescence was roughly half that of what was measured based on gel band intensity (2).

We emphasize that calibration values are specific to the aptamer and solution conditions used for the experiment and can't be applied universally. In fact, even drawing conclusions from relative comparisons when aptamer or solution conditions are modified is not advised since they are likely to alter the relationship between amount of RNA and fluorescence. Many other fluorescent aptamer sequences have been described and designed, improving on the original Spinach and Spinach-mini sequence used here, by increasing thermostability, folding propensity, quantum-yields, etc. (4–10). As a result, the same amount of transcript generated containing a different aptamer sequence, will invariably yield a different fluorescent signal. Additionally, each aptamer sequence should be tested via gel assays **(Supplementary Figure 8)** to ensure the sequence has no effect on transcription kinetics.

Similarly, care must be taken if one is interested in the effects of solution conditions on transcription. For instance, salt identity and concentration can have large effects on both transcription kinetics (11–13) and aptamer folding/dye binding (8, 14, 15). As a result, without performing a calibration curve at each unique salt-concentration tested, one could not comment on whether the change in fluorescence is due to transcriptional activity, aptamer folding, or some combination of the two. For this reason, we strongly suggest keeping conditions that affect RNA folding (salt, $Mg^{2+}$, pH, temperature, DMSO, etc.) constant when comparing effects of transcription factors or promoter sequence, or when performing a titration series. For instance, if titrating a component stored in a different buffer than that of the reaction buffer, use the same volume percentage of the component and its storage buffer for each condition, or dialyze all reaction components into the same buffer.

### Supplementary Tables

**Table S1: Insert sequences for plasmid DNA constructs.** The non-template strand sequences displayed below are the requested inserts that are cloned into the pTwist, High Copy, $Amp^R$ plasmid vector (Twist Bioscience). Sequence numbering is relative to the transcription start site (blue, numbered as +1,) where the genomic locations are provided based on the *Mtb* H37Rv numbering**.** The terminator corresponds to 80 nt of the *E. coli rrnB*P1 $T_1$ genomic sequence (red, 6608 to 6687 based on numbering in Ref. (16)). The aptamer corresponds to the Spinach-mini RNA sequence (green) (17). Non-genomic-derived sequences were included as Forward (F-) and Reverse (R-) primer annealing sites (yellow) and can be used to amplify the sequence insert from the plasmid to make linear templates.

| | |
|---|---|
| **1. *Mtb rrnA*P3** | |
| **Description** | F-primer *rrnA*P3 aptamer +39bp *rrnA*P3 +80bp terminator R-primer |
| **DNA Sequence (Non-Template) (5'-3')** | GAGCTCGGTACCCGGGGATCATCTATGGATGACCGAACCTGGTCTTGACTCCATTGCCGGATTTGTA<br>TTAGACTGGCAGGGTTGCCCCGAAGCGGGCGGAAACAAGCAAGCGACGCGACCGAAATGGTGAAG<br>GACGGGTCCAGTGCTTCGGCACTGTTGAGTAGAGTGTGAGCTCCGTAACTGGTCGCGTCGTGTTGTT<br>TGAGAACTCAATAGTGTGTTTGGTGGTTTCACCAGGCATCAAATAAAACGAAAGGCTCAGTCGAAAGA<br>CTGGGCCTTTCGTTTTATCTGTTGTTTGTCGGTGAACGCTCTCGCCTCTCCCCGCGCGTTGGC |
| **Genomic Location** | 1,471,597 (-60) to 1,471,687 (+31); 1,471,688(+32) to 1,471,726(+70) |
| **2. *Mtb rrnA*P3 without aptamer** | |
| **Description** | F-primer *rrnA*P3 +80bp terminator R-primer random |
| **DNA Sequence (Non-Template) (5'-3')** | GAGCTCGGTACCCGGGGATCATCTATGGATGACCGAACCTGGTCTTGACTCCATTGCCGGATTTGTA<br>TTAGACTGGCAGGGTTGCCCCGAAGCGGGCGGAAACAAGCAAGCGTGTTGTTTGAGAACTCAATAG<br>TGTGTTTGGTGGTTTCACCAGGCATCAAATAAAACGAAAGGCTCAGTCGAAAGACTGGGCCTTTCGTT<br>TTATCTGTTGTTTGTCGGTGAACGCTCTCGCCTCTCCCCGCGCGTTGGCCCCGTCACCAGATTAAGC<br>GGAAAGCGCAAAGCAAGGATTGGGCGTTCCGC |
| **Genomic Location** | 1,471,597 (-60) to 1,471,726(+70) |
| **3. PromoterLESS** | |
| **Description** | F-primer *rrnA*P3 (Δ -49 to -1 of *rrnA*P3) *rrnA*P3 aptamer +39bp *rrnA*P3 +80bp terminator R-primer |
| **DNA Sequence (Non-Template) (5'-3')** | GAGCTCGGTACCCGGGGATCATCTATGGATCATGTTTGACAGCTTATCATCGGAGCTCTCGAGTCTA<br>GAATCGATCCCCGGTTGCCCCGAAGCGGGCGGAAACAAGCAAGCGACGCGACCGAAATGGTGAAG<br>GACGGGTCCAGTGCTTCGGCACTGTTGAGTAGAGTGTGAGCTCCGTAACTGGTCGCGTCGTGTTGTT<br>TGAGAACTCAATAGTGTGTTTGGTGGTTTCACCAGGCATCAAATAAAACGAAAGGCTCAGTCGAAAGA<br>CTGGGCCTTTCGTTTTATCTGTTGTTTGTCGGTGAACGCTCTCGCCTCTCCCCGCGCGTTGGC |
| **Genomic Location** | 1,471,597 (-60) to 1,471,606 (-50); 1,471,657 (+1) to 1,471,687 (+31); 1,471,688(+32) to 1,471,726(+70) |

#### *Descriptions*

1. ***Mtb rrnA*P3**: Primary template used in this work. Contains both the Spinach-mini aptamer and the *E. coli rrnB*P1 $T_1$ terminator. The *Mtb rrnA*P3 genomic sequence (purple) is present both upstream (from –60 to +31) and downstream (from +32 to +70) of the aptamer.
2. ***Mtb rrnA*P3 without aptamer**: Same design as construct #1, except the aptamer sequence was removed. The randomized sequence (cyan) added after the terminator and R-primer was needed to reach the required minimal insert length.
3. **"PromoterLESS":** Same design as construct #1 except that –49 to –1 of the *Mtb rrnA*P3 genomic DNA was replaced with a randomized sequence (orange).

**Table S2: Primer and final sequences for linear DNA constructs.** Note that the genomic sequence numbering and color coding below follows the same format as described in **Supplementary Table 1**. Underlined sequences represent those within the 150 bp annealing region. An overview of the preparation steps is included: Following annealing and extension of the 200 nt oligonucleotide primers (sequences shown under blue heading), a 250 bp construct is obtained (final non-template sequence shown under grey heading). Once extended, the genomic *rrnA*P3 sequence from –60 to +31 is present upstream of the aptamer and from +32 to +70 is present downstream of the aptamer, as in the DNA plasmid constructs. The PCR template can then be amplified the template using the primer sequences shown in yellow. We added a biotin molecule to the Forward, 20 nt primer, attached to the 5'-end via a standard C6 spacer (code for modification is /5Biosg/ when ordering from Integrated DNA Technologies, Inc.). The biotin addition was in hopes to reduce end-binding/initiation effects.

**Overview of preparing 250 bp linear constructs**

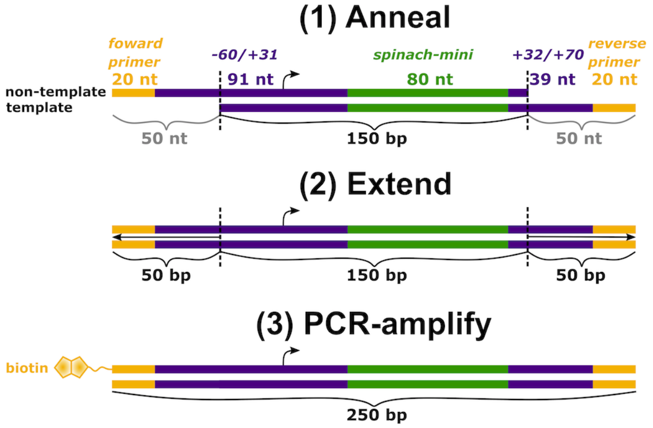


**Primers for annealing/extending**

| DNA Sequence (5'-3') | Genomic Location | Strand | Length (nt) |
|---|---|---|---|
| GAGCTCGGTACCCGGGGATCATCTATGGATGACCGAACCTGGTCTTGACT<br>CCATTGCCGGATTTGTATTAGACTGGCAGGGTTGCCCCGAAGCGGGCGGA<br>AACAAGCAAGCGACGCGACCGAAATGGTGAAGGACGGGTCCAGTGCTTCG<br>GCACTGTTGAGTAGAGTGTGAGCTCCGTAACTGGTCGCGTCGTGTTGTTT | 1,471,597 (-60) to 1,471,687 (+31); 1,471,688 (+32) to 1,471,696 (+41) | Non-template | 200 |
| GCCAACGCGCGGGGAGAGGCTGAAACCACCAAACACACTATTGAGTTCTC<br>AAACAACACGACGCGACCAGTTACGGAGCTCACACTCTACTCAACAGTGCC<br>GAAGCACTGGACCCGTCCTTCACCATTTCGGTCGCGTCGCTTGCTTGTTTC<br>CGCCCGCTTCGGGGCAACCCTGCCAGTCTAATACAAATCCGGCAATGG | 1,471,726 (+70) to 1,471,688 (+32); 1,471,687 (+31) to 1,471,627 (-30) | template | 200 |

**Final non-template sequence**

| | |
|---|---|
| Description | F-primer *rrnA*P3 aptamer +39bp *rrnA*P3 R-primer |
| DNA Sequence (Non-Template) (5'-3') | GAGCTCGGTACCCGGGGATCATCTATGGATGACCGAACCTGGTCTTGACTCCATTGCCGGATTT<br>GTATTAGACTGGCAGGGTTGCCCCGAAGCGGGCGGAAACAAGCAAGCGACGCGACCGAAATG<br>GTGAAGGACGGGTCCAGTGCTTCGGCACTGTTGAGTAGAGTGTGAGCTCCGTAACTGGTCGCG<br>TCGTGTTGTTTGAGAACTCAATAGTGTGTTTGGTGGTTTCAGCCTCTCCCCGCGCGTTGGC |
| Genomic Location | 1,471,597(-60) to 1,471,687(+31); 1,471,688(+32) to 1,471,726(+70) |

**Table S3: Calculated $K_m$, $V_{max}$ and *n* values from NTP titrations.** Values and errors provided are the results from fits to the averaged data (see "Statistical Analyses" section in Methods for more details). For titrations in the context of plasmid DNA templates a conversion factor of 58.8 fluorescence units/ 1 nM RNA concentration was used, based on the calibration in **Figure 5D**. All NTP titrations were performed with 100 nM *Mtb* RNAP and either 5 nM plasmid or 25 nM linear *Mtb* rrnAP3 DNA.

| | **Plasmid DNA** | | | **Linear DNA** | | |
|---|---|---|---|---|---|---|
| **Titrated NTP** | $K_{m,app}$ (μM) | $V_{max}$ (nM RNA/min) | ***n*** | $K_{m,app}$ (μM) | $V_{max}$ (AU/min) | ***n*** |
| **ALL** | 44 ± 5 | 2.8 ± 0.2 | 2.1 ± 0.4 | 138 ± 30 | 5.6 ± 0.3 | 2.1 ± 0.6 |
| **GTP** | 16 ± 2 | 2.6 ± 0.2 | 1.9 ± 0.4 | 44 ± 3 | 5.6 ± 0.3 | 1.3 ± 0.1 |
| **UTP** | 5.8 ± 1.0 | 2.9 ± 0.2 | 1.2 ± 0.1 | 12 ± 3 | 5.6 ± 0.2 | N/A |
| **CTP** | 2.2 ± 1.5 | 2.6 ± 0.1 | N/A | 1.3 ± 0.2 | 5.6 ± 0.1 | N/A |
| **ATP** | 0.8 ± 0.3 | 3.1 ± 0.3 | N/A | 1.4 ± 1 | 5.7 ± 0.3 | N/A |

* N/A = not applicable, as **Eq. 1** was used for fitting which lacked the additional parameter *n*

### Supplementary Figures

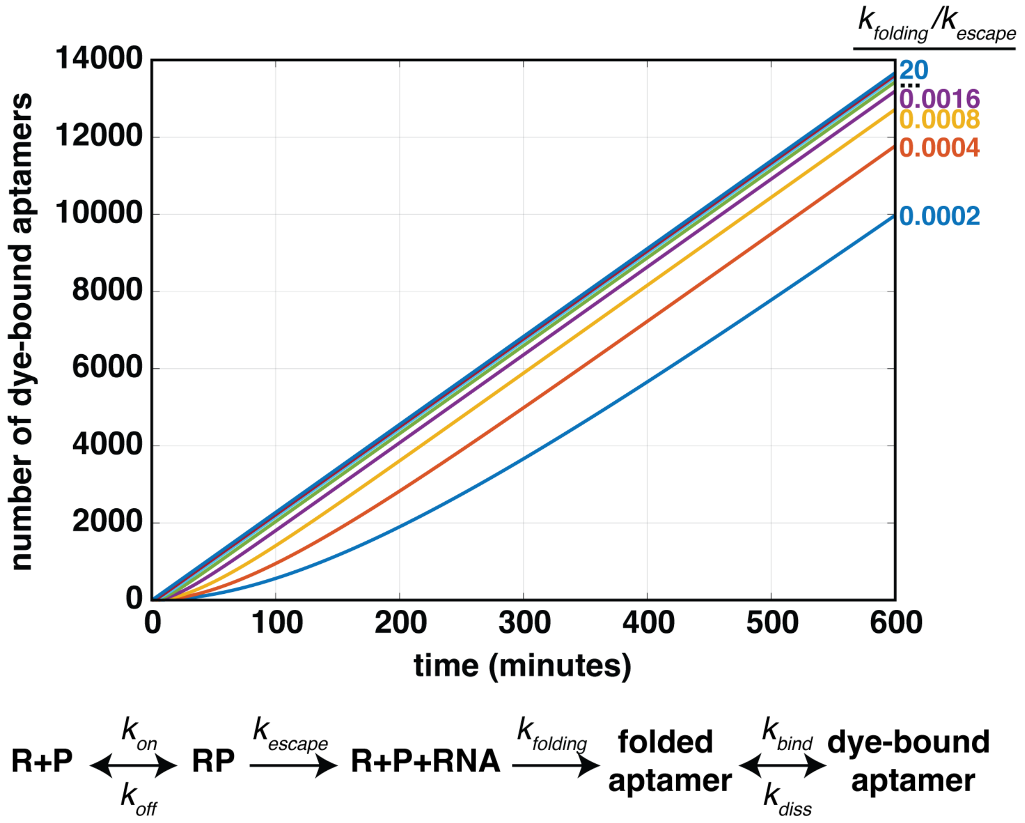


**Figure S1: Simulations where aptamer-folding is rate-limiting.** Simulations of transcription and fluorescence detection were performed according to the kinetic scheme above (Methods), where R is RNAP and P is the promoter template. To assess how a slow molecular process downstream of the promoter, such as aptamer folding ($k_{folding}$), may affect the steady-state rate of initiation, the rate of folding was titrated from 0.0002x to 20x the rate of promoter escape. The resulting curves illustrate that the steady-state rate of dye-bound aptamers (linear slope) is independent of the aptamer folding rate. However, when $k_{\mathrm{folding}}$ is slower than $k_{\mathrm{escape}}$, an increase in the time to reach steady-state (i.e., a lag-time) is observed. We note that in the context of this model, we don't explicitly model elongation, but it can be considered to be included in the aptamer folding rate as multiple elongation reactions can take place in parallel on a single template. In addition, previous studies illustrated the insertion of up to 500 bp in sequence between the promoter and the aptamer sequence had no effect on the observed lag-time (3). The parameters used in the simulation were as follows: $k_{\mathrm{on}}$ = 0.01 $nM^{-1}s^{-1}$, $k_{\mathrm{off}}$ = 0.1 $s^{-1}$, $k_{\mathrm{escape}}$ = 0.5 $s^{-1}$, $k_{\mathrm{folding}}$ = (0.0001 – 10 $s^{-1}$), $k_{\mathrm{bind}}$ = 0.00004 $nM^{-1}s^{-1}$, $k_{\mathrm{diss}}$ = 0.001 $s^{-1}$, [RNAP] = 200 nM, [dye] = 20 μM.

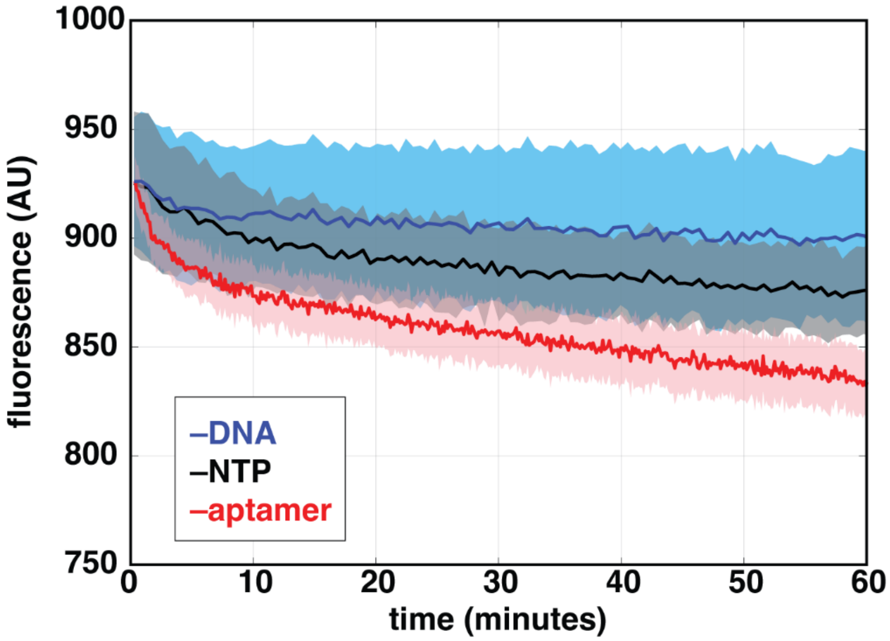


**Figure S2: Real-time fluorescent signal in the absence of aptamer formation.** Fluorescent signal was monitored as a function of time with 100 nM *Mtb* RNAP and 20 μM DFHBI under conditions where aptamer formation was prevented, either by leaving out 500 μM all NTPs (black), 5 nM *rrnA*P3 plasmid DNA (blue), or including the plasmid DNA template which lacked the aptamer sequence (red). Leaving out any one of these reagents results in no increase in fluorescence, but rather only a slow decay. As a result, in each data acquisition, a negative control is collected and subtracted from the experimental data to correct for this decay. These curves are similar, but there can be small variations of the shape and amplitude of the decay. For this reason, we always include a negative control to be used to correct the data collected in the same experiment. While this correction doesn't significantly affect moderate to high signal experiments (i.e., it only represents ~1% of the signal change observed at saturating NTPs), it becomes crucial when studying systems with low signals.

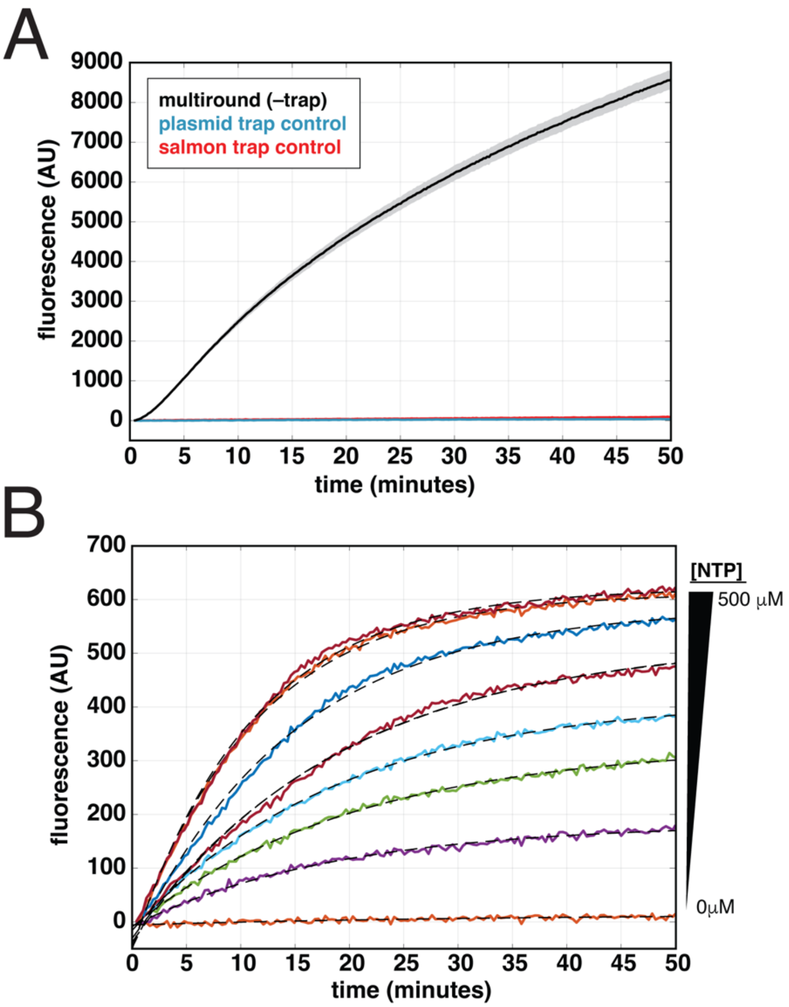


**Figure S3: Single-round trap controls and kinetics. (A)** Incubation of either 75 μg/mL of salmon-sperm DNA (red) or 150 nM "without aptamer" plasmid (blue) trap prior to the addition of NTPs results in the complete abrogation of fluorescent signal, compared to conditions of 100 nM *Mtb* RNAP, 5 nM *Mtb rrnA*P3 plasmid DNA and 500 μM all NTPs without trap pre-incubation (black). **(B)** Titration of all NTPs under single-round conditions, where pre-incubated RNAP-promoter complexes (100 nM *Mtb* RNAP, 5 nM *Mtb rrnA*P3 plasmid DNA) were initiated with various concentrations of all NTPs and saturating amounts of salmon-sperm DNA. Dashed lines represent single-exponential fits at each NTP concentration tested. Here, fluorescence amplitude increases with increasing NTP concentration, as expected for a system that is prone to dissociation during promoter escape at low NTP concentrations.

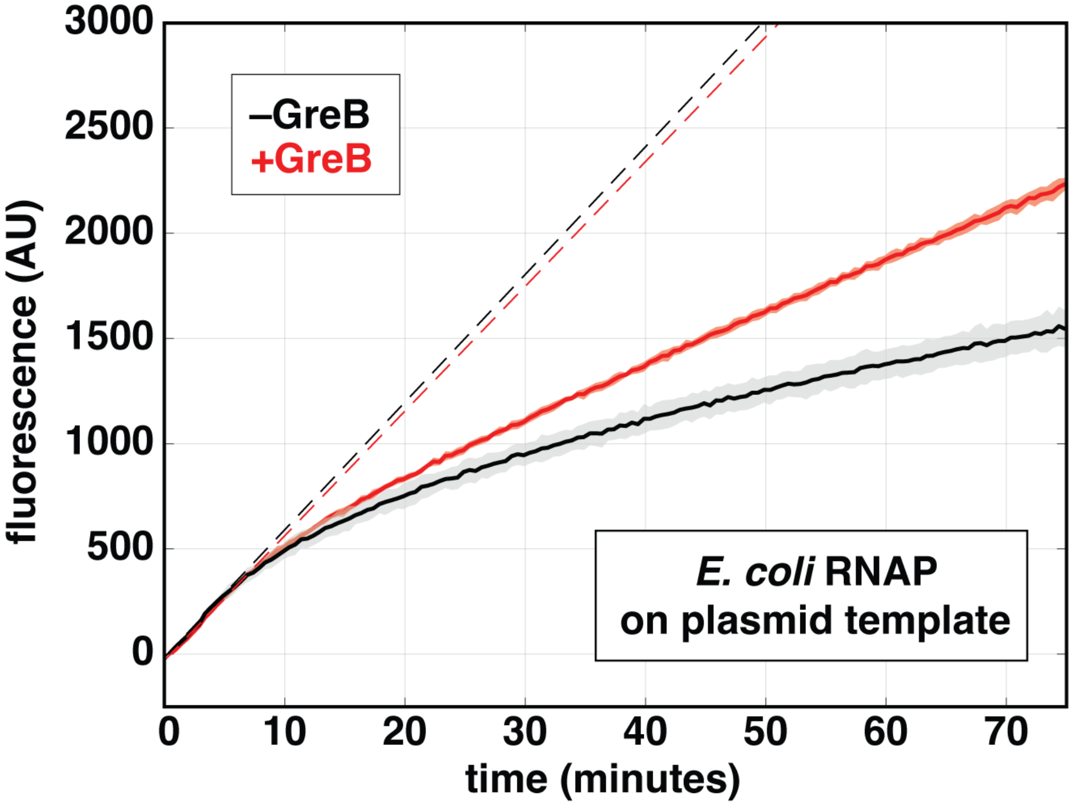


**Figure S4: GreB affects the long-term behavior of fluorescent traces.** Experiments were performed using 100 nM *E. coli* $\sigma^{70}$ RNAP, 500 µM all NTPs, 5 nM *Mtb rrnA*P3 plasmid DNA in the presence and absence of 1 µM *E. coli* GreB. Dotted lines represent the fits to the initial steady-state regime occurring from 0 – 10 minutes. The effect of GreB can be seen at longer time-scales past the initial steady-state.

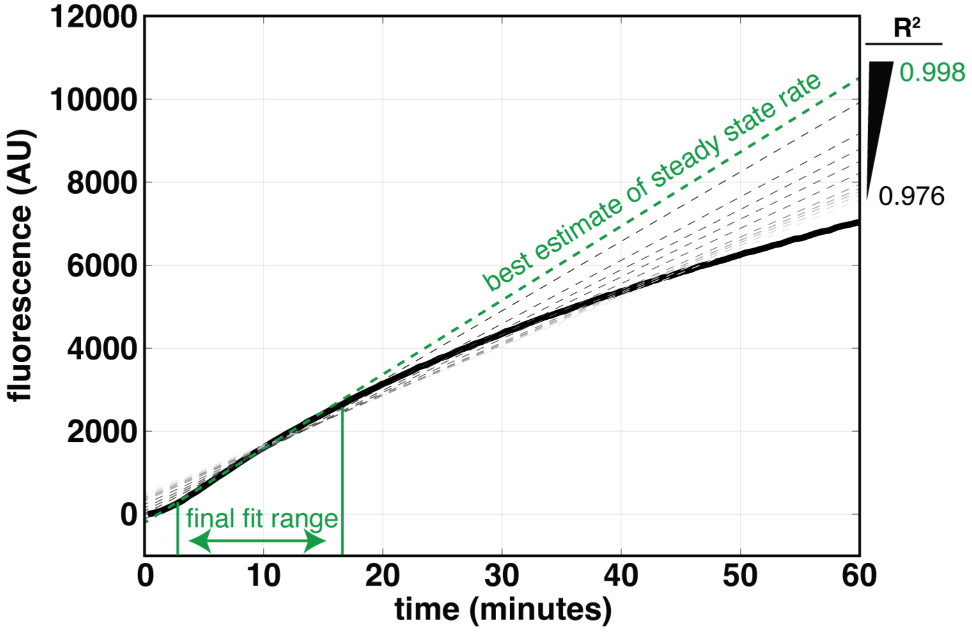


**Figure S5: Overview of variable-time, iterative linear fitting approach.** Examples of recursive linear fitting using our code (https://github.com/egalburt/aptamer-flux-fitting) to illustrate how goodness of fit can change depending on the time interval. With each iteration, a shorter segment of the data is fit, and both the measured rate and $R^2$ value increase until a threshold $R^2$ is reached (in this case, 0.998).

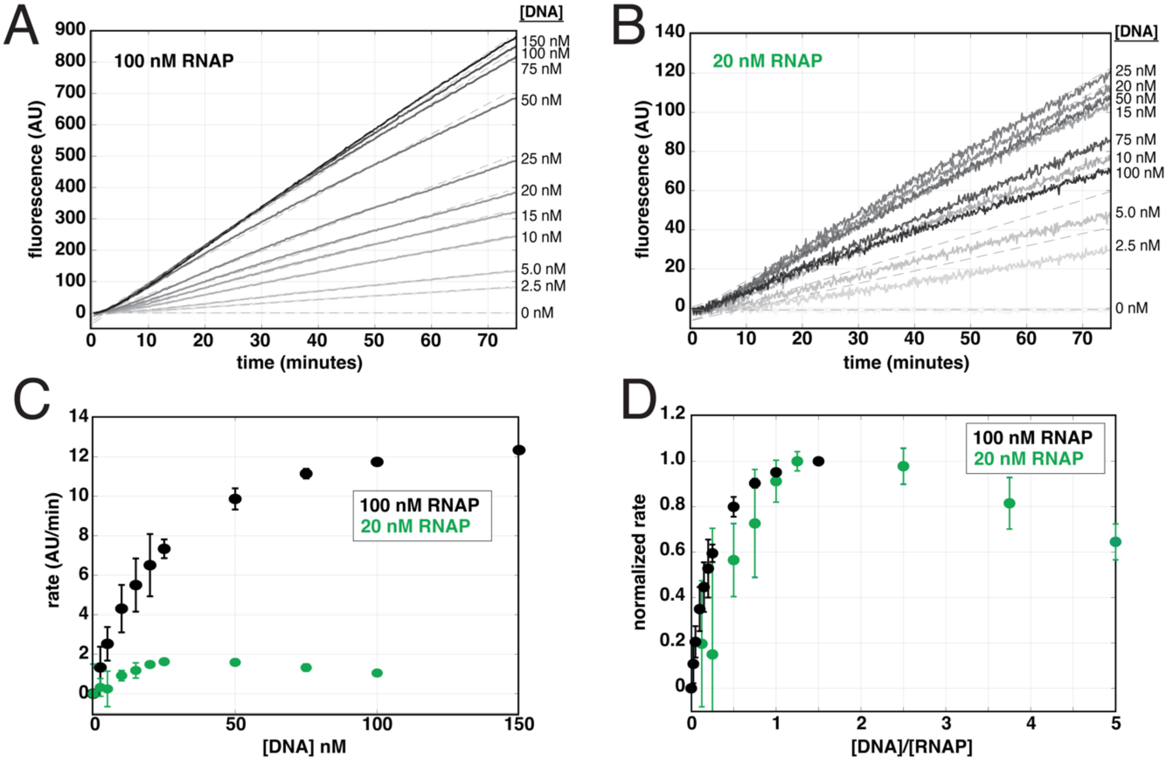


**Figure S6: Real-time data for DNA titrations on linear templates.** Compare to plasmid data in **Figure 4** of the main text. Data was obtained at 500 µM all NTPs, titrating *Mtb rrnA*P3 linear DNA template (2.5 – 150 nM) at either **(A)** 100 nM or **(B)** 20 nM *Mtb* RNAP concentrations**.** The unbiased linear fits of the early times are shown in grey dotted lines for each trace. **(C)** Steady-state rates obtained from the linear fits in (A) and (B) for 20 nM (green) and 100 nM (black) RNAP, plotted as a function of *rrnA*P3 linear DNA concentration. **(D)** Steady-state rates, normalized from zero to one based on the lowest and highest rate obtained at each RNAP concentration, plotted as a function of the ratio of [DNA]:[RNAP] concentrations.

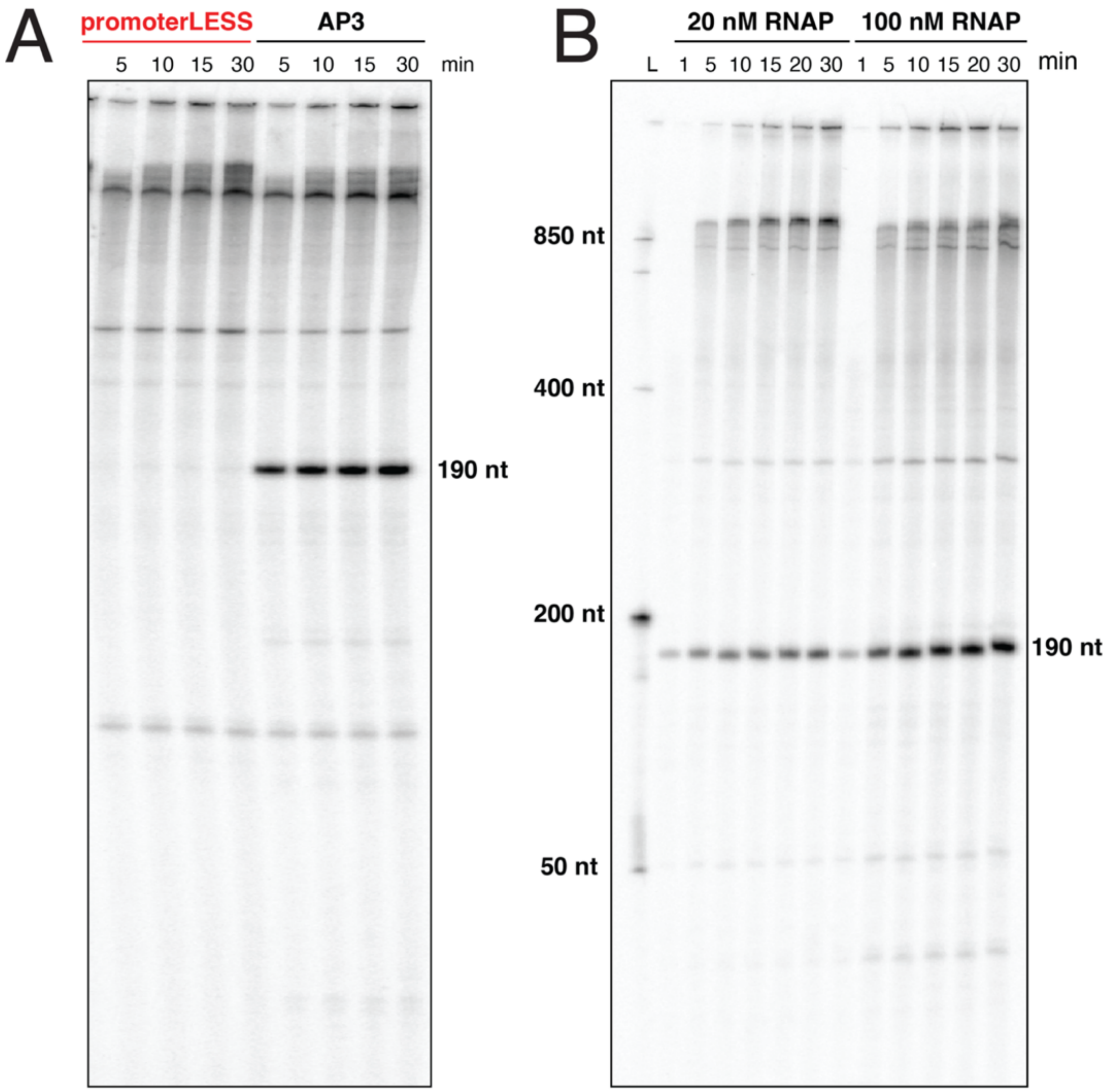


**Figure S7: Full gel images of the data presented in main text Figure 5. (A)** Time-dependent monitoring of all transcription bands observed with the *rrnA*P3 and promoterLESS plasmids and **(B)** with the *rrnA*P3 plasmid at 20 and 100 nM RNAP concentrations. All reactions were performed with *Mtb* RNAP, 5 nM plasmid DNA templates, 500 μM all NTPs and 20 μM DFHBI. The promoter-derived band containing the aptamer sequence runs to a length of ~ 190 nt.

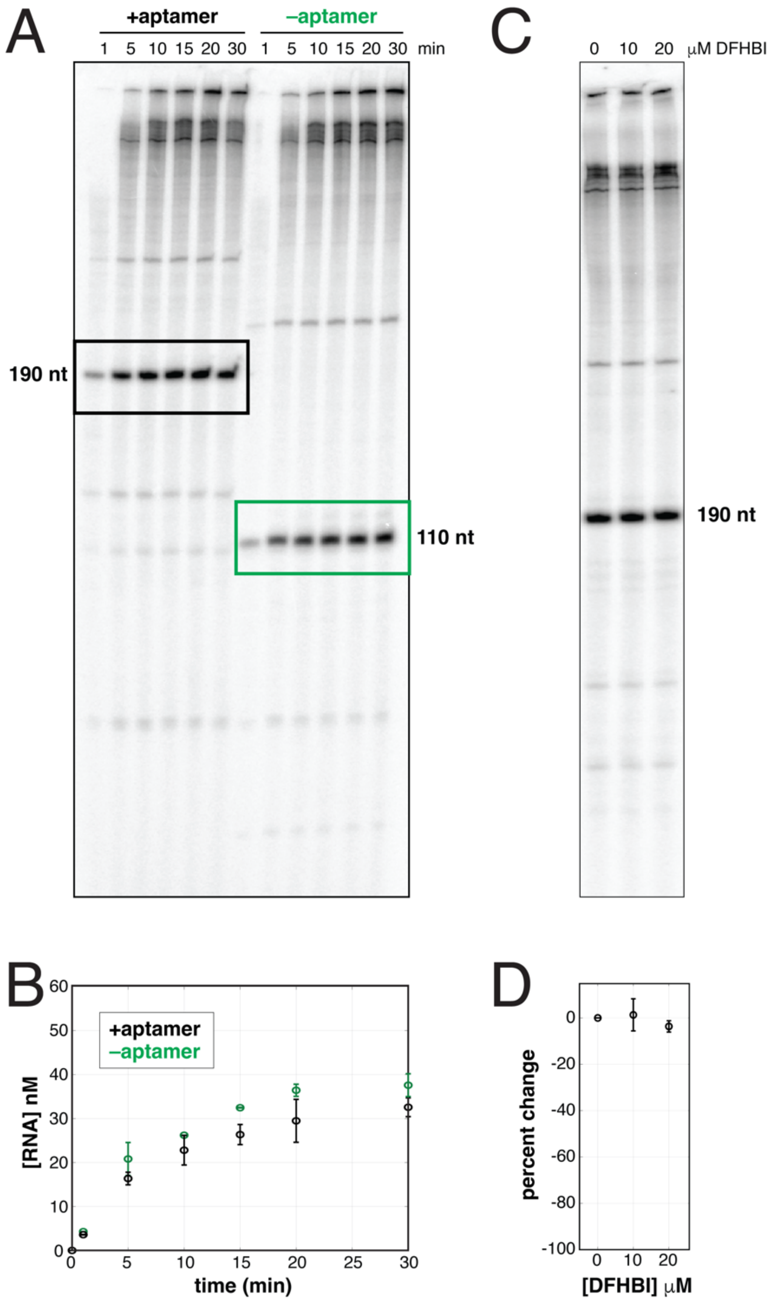


**Figure S8: Gel-based analysis of the effect of aptamer sequence and DFHBI dye. (A)** Time-dependent monitoring of all transcription bands observed with the *rrnA*P3 plasmids with and without the aptamer. **(B)** Band quantification in units of nM RNA of the promoter-derived product in (A) plotted as a function of time. **(C)** Measurement of all transcription bands observed with the *rrnA*P3 plasmid post 30 min after reaction initiation as a function of DFHBI concentration. **(D)** Band quantification of the promoter-derived product in (C) normalized to the signal obtained with 0 μM dye. Note that in (A) the promoter-derived band from the no-aptamer plasmid runs lower, as that 80 nt sequence was removed (**Supplementary Table 1**).

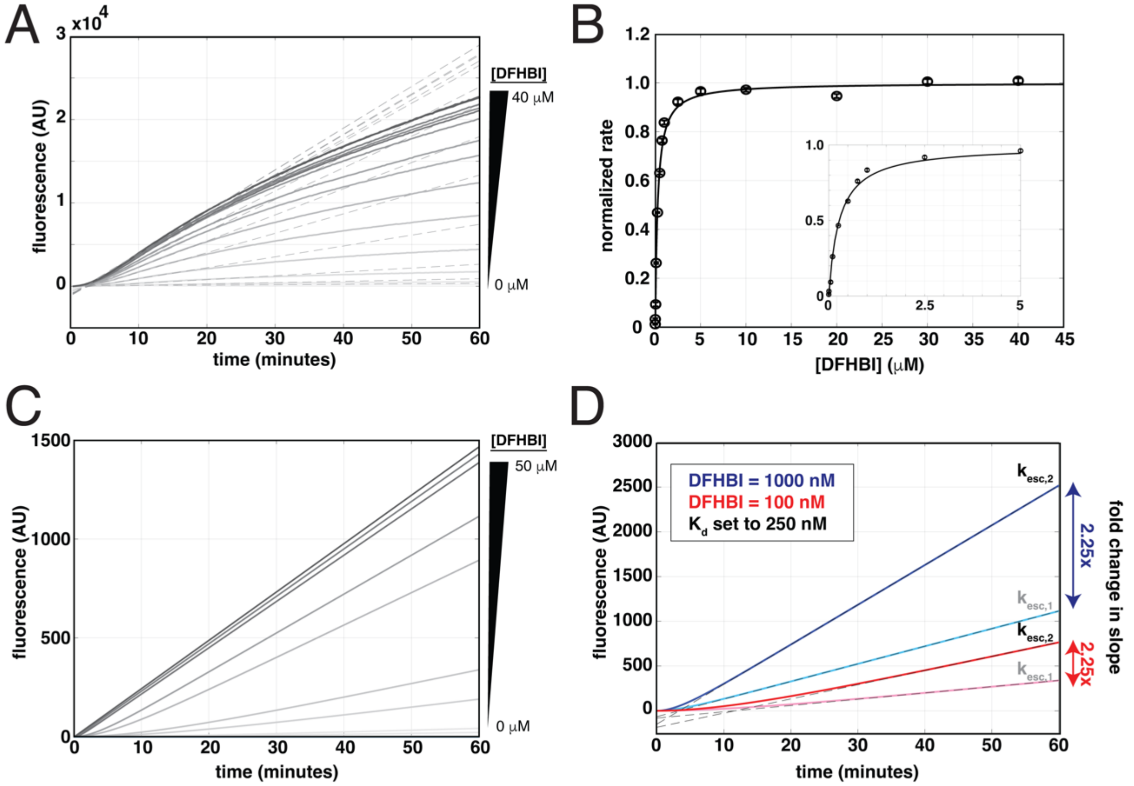


**Figure S9: Concentration dependencies of DFHBI on the fluorescent-aptamer signal. (A)** DFHBI dye titration with 100 nM *Mtb* RNAP, 500 µM NTPs, and 5 nM *rrnA*P3 plasmid. The unbiased linear fits of the early times are shown in grey dotted lines for each trace. **(B)** Normalized steady-state rates plotted as a function of DFHBI concentration. Fit to a hyperbolic function yielded a midpoint of 270 nM, comparable to the binding affinities measured to Spinach, Spinach2, and Broccoli (5, 7, 17). Inset in (B) depicts the titration, from 0 – 5 µM dye concentrations. The steady-state rates measured in this regime change as the fraction of transcribed aptamer that is bound by dye changes. However, the underlying rate of transcription does not change as can be seen via the gel-based experiments in **Supplementary Figure S8C,D**. In essence then, dye concentration simply allows one to titrate the gain on the signal. **(C)** Kinetic simulations were performed using the model in **Supplementary Figure 1** with fixed rate constants and varying concentrations of DFHBI dye (Methods). As seen in our experimental data, increasing dye concentration increases the slope of the curves until saturating at higher concentrations. **(D)** A comparison of the same kinetic scheme with two different escape rates ($k_{escape,1}$ = 3X $k_{escape,2}$) in the presence of two different dye concentrations (cyan/blue curves and pink/red curves). Although the absolute slopes of the curves at higher dye concentration are steeper, the fold change in the slopes between the two kinetic schemes remains the same. This illustrates the importance of making comparisons using the same dye concentrations, but also demonstrates the independence of mechanistic conclusions regarding initiation from the absolute concentration of dye.

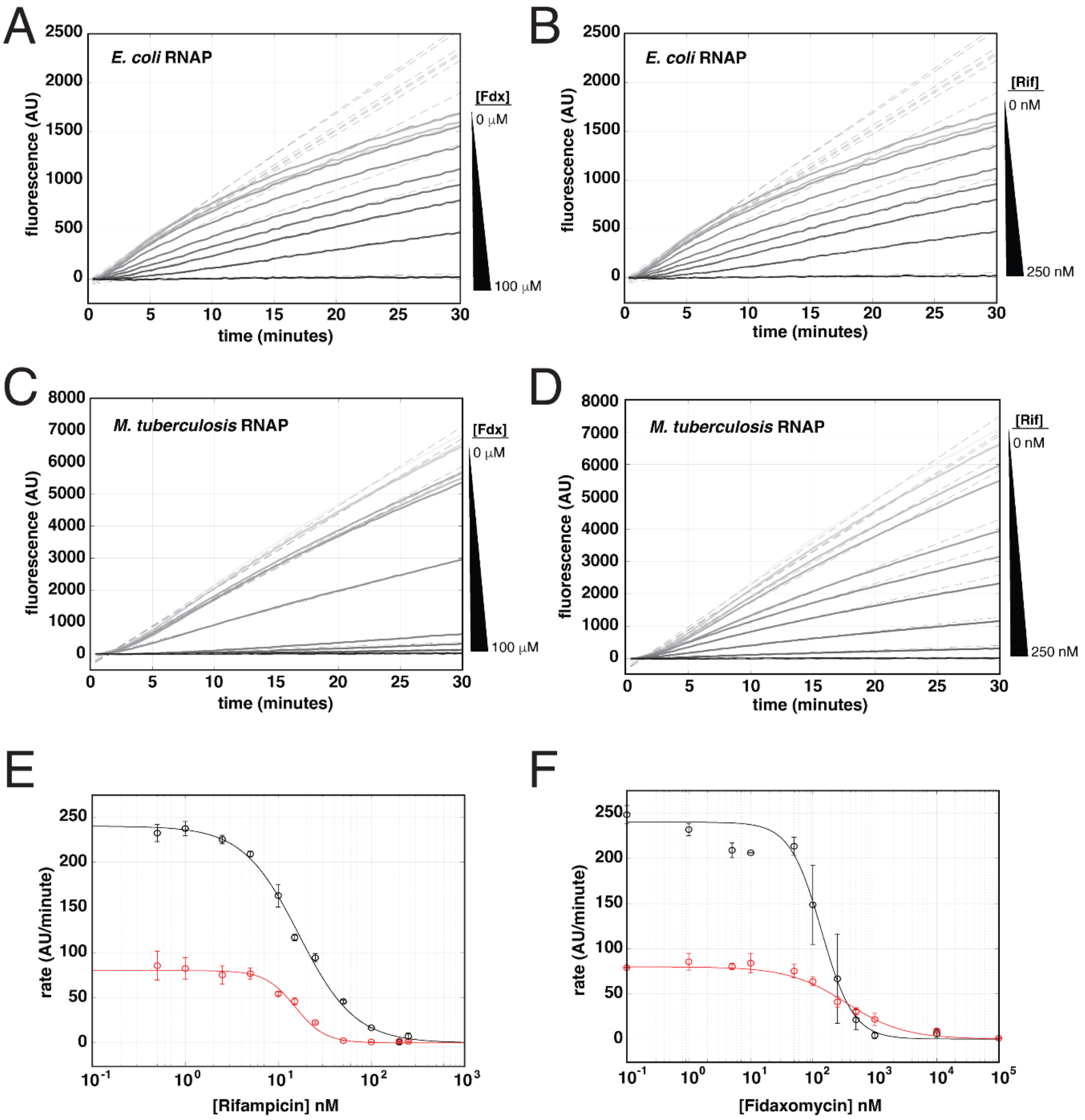


**Supplementary Figure S10: Real-time data for antibiotic titrations and unnormalized $IC_{50}$ fits.** Real-time data for Fidaxomicin and Rifampicin titrations (Methods) shown for *E. coli* (**A**, **B**) and *Mtb* (**C**, **D**) RNAPs. Dotted lines in (A-D) represent linear fits to the steady-state regime. The resulting steady-state rates are plotted as a function of Rifampicin (**E**) and Fidaxomicin (**F**) concentration for *Mtb* (black) and *E. coli* (red) RNAPs and are fit to **Eq. 3** for a measure of the respective $IC_{50}$s. Data in **Figure 8** is normalized from one to zero based on the fits presented in (E) and (F).
